# The CRP-like transcriptional regulator MrpC curbs c-di-GMP and 3’, 3’ cGAMP nucleotide levels during development in *Myxococcus xanthus*

**DOI:** 10.1101/2021.10.14.464357

**Authors:** Sofya Kuzmich, Patrick Blumenkamp, Doreen Meier, Alexander Goesmann, Anke Becker, Lotte Søgaard-Andersen

## Abstract

*Myxococcus xanthus* has a nutrient-regulated biphasic lifecycle forming predatory swarms in the presence of nutrients and spore-filled fruiting bodies in the absence of nutrients. The second messenger c-di-GMP is essential during both stages of the lifecycle; however, different enzymes involved in c-di-GMP synthesis and degradation as well as several c-di-GMP receptors are important during distinct lifecycle stages. To address this stage specificity, we determined transcript levels using RNA-seq and transcription start sites using Cappable-seq during growth and development at a genome-wide scale. All 70 genes encoding c-di-GMP associated proteins were expressed, with 28 up-regulated and 10 down-regulated during development. In particular, the three genes encoding enzymatically active proteins with a stage-specific function were expressed stage-specifically. By combining operon mapping with published ChIP-seq data for MrpC (Robinson et al., 2014), the CRP-like master regulator of development, we identified nine developmentally regulated genes as regulated by MrpC. In particular, MrpC directly represses expression of *dmxB*, which encodes the diguanylate cyclase DmxB that is essential for development and responsible for the c-di-GMP increase during development. Moreover, MrpC directly activates transcription of *pmxA*, which encodes a bifunctional phosphodiesterase that degrades c-di-GMP and 3’, 3’ cGAMP *in vitro* and is essential for development. Thereby, MrpC regulates and curbs the cellular pools of c-di-GMP and 3’, 3’ cGAMP during development. We conclude that temporal regulation of the synthesis of proteins involved in c-di-GMP metabolism contributes to c-di-GMP signaling specificity. MrpC is important for this regulation, thereby being a key regulator of developmental cyclic di-nucleotide metabolism in *M. xanthus*.

**Importance:** The second messenger c-di-GMP is important during both stages of the nutrient-regulated biphasic lifecycle of *Myxococcus xanthus* with the formation of predatory swarms in the presence of nutrients and spore-filled fruiting bodies in the absence of nutrients. However, different enzymes involved in c-di-GMP synthesis and degradation are important during distinct lifecycle stages. Here, we show that the three genes encoding enzymatically active proteins with a stage-specific function are expressed stage-specifically. Moreover, we find that the master transcriptional regulator of development MrpC directly regulates expression of *dmxB*, which encodes the diguanylate cyclase DmxB that is essential for development, and of *pmxA*, which encodes a bifunctional phosphodiesterase that degrades c-di-GMP and 3’, 3’ cGAMP *in vitro* and is essential for development. We conclude that temporal regulation of the synthesis of proteins involved in c-di-GMP metabolism contributes to c-di-GMP signaling specificity, and that MrpC plays an important role in this regulation.

## Introduction

In bacteria, signaling by nucleotide-based second messengers have important functions in adaptive responses to environmental changes (1-6). 3’-5’, 3’-5 cyclic di-GMP (c-di-GMP) is a versatile second messenger that regulates numerous processes including exopolysaccharide (EPS) synthesis, biofilm formation, cell cycle progression, virulence, motility, and multicellular development (1, 2). c-di-GMP is synthesized by diguanylate cyclases (DGCs), which contain the conserved GGDEF domain, and degraded by phosphodiesterases (PDEs), which contain an EAL or HD-GYP domain (1, 2). The effects of changing c-di-GMP levels are implemented by c-di-GMP binding receptors, which regulate downstream responses at the transcriptional, translational or post-translational level (1, 2). Reflecting the versatility of c-di-GMP, c-di-GMP receptors comprise a variety of proteins with little sequence homology, including enzymatically inactive DGC and EAL domain proteins (7-11), PilZ-domain proteins (12-16), MshEN-domain proteins (17, 18), and proteins of different transcription factor families (19-26). Among these receptors, enzymatically inactive DGC and EAL domain proteins as well as PilZ- and MshEN domains can be bioinformatically predicted (17, 27).

Often individual bacterial genomes encode multiple DGCs, PDEs and c-di-GMP receptors (1). Yet, inactivation of individual genes for DGCs, PDEs and c-di-GMP receptors can result in distinct phenotypes, underscoring that specific signaling modules exist. Thus, a central question is how this signaling specificity is accomplished. Three mutually non-exclusive models have been proposed to explain this specificity (1, 28, 29). Firstly, individual signaling modules can be temporally separated based on differential regulation of their synthesis and/or degradation; secondly, individual signaling modules can be spatially separated by protein complex formation or by localizing to distinct subcellular locations; and, thirdly, effectors of different signaling modules may have different binding affinities for c-di-GMP.

*Myxococcus xanthus* is a model organism for studying social behaviors and cell differentiation in bacteria (30). *M. xanthus* has a nutrient-regulated biphasic life cycle. In the presence of nutrients, cells form predatory swarms that spread coordinately using type IV pilus (T4P)-dependent motility and gliding motility (31, 32). Upon nutrient depletion, *M. xanthus* initiates a developmental program that culminates in the formation of multicellular, spore-filled fruiting bodies while cells that remain outside fruiting bodies differentiate to either so-called peripheral rods or undergo cell lysis (33-35). Nucleotide-based second messengers have important roles during both stages of the lifecycle: During growth, c-di-GMP is important for type IV pili-dependent motility and for regulation of motility (36, 37). During development, the starvation-induced activation of the stringent response with synthesis of the second messenger (p)ppGpp is required and sufficient to initiate development (38, 39). Moreover, the cellular c-di-GMP level increases dramatically during development, and this increase is essential for completion of development (40). Development also depends on global transcriptional changes (41), regulation of motility (31, 32) and cell-cell signaling (30, 42).

Several transcription factors that are essential for fruiting body formation and sporulation have been identified (41). Among these, MrpC is a member of the cAMP receptor protein (CRP) family of transcription factors (43) and has been referred to as a master regulator of development (41). Currently, no ligand for MrpC has been reported, and MrpC on its own binds target promoters *in vitro* (44-51). MrpC alone is a negative autoregulator (44) and directly activates transcription of *fruA* (45), which encodes a transcriptional regulator that is also essential for development (52, 53). MrpC and FruA jointly regulate the expression of multiple genes during development (46-51).

Systematic inactivation of all 36 genes for GGDEF-domain proteins, EAL-domain proteins, and HD-GYP-domain proteins identified only three enzymatically active proteins that are important during growth and development under standard laboratory conditions. Interestingly, each of the three proteins are important during a distinct stage of the lifecycle. The DGC DmxA is important for T4P-dependent motility in the presence of nutrients but not for development (37, 40). By contrast, the DGC DmxB and the HD-GYP-type PDE PmxA are exclusively important for development (37, 40). DmxB is the DGC responsible for the dramatic increase in the c-di-GMP level during development (40). PmxA degrades c-di-GMP as well as the di-nucleotide 3’-5’, 3’-5’ cyclic GMP-AMP (cGAMP) *in vitro* and with the highest activity towards cGAMP (40, 54). Lack of PmxA does not lead to significant changes in the c-di-GMP level during development (40) while it remains unknown how lack of PmxA may affect cGAMP accumulation *in vivo*.

Several c-di-GMP receptors have been experimentally verified in *M. xanthus*. The histidine protein kinase SgmT contains an enzymatically inactive GGDEF domain that binds c-di-GMP and works together with the DNA binding response regulator DigR to regulate extracellular matrix composition during growth and development (8, 37). The enhancer binding protein Nla24 also binds c-di-GMP and is important for motility during growth as well as development (40, 55, 56). Systematic inactivation of all 24 genes encoding PilZ-domain proteins identified PixA and PixB as c-di-GMP receptors that regulate motility (36). While PixA is important only during growth, PixB is crucial during growth and development (36). Finally, the ribbon-helix-helix proteins CdbA and CdbB bind c-di-GMP (57). CdbA is an essential nucleoid-associated protein important for chromosome organization and segregation (57).

With the exception of DmxB, synthesis of which is strongly up-regulated during development (40), it is not understood how c-di-GMP metabolizing enzymes and some verified receptors are functionally restricted to either growth or development. To increase our understanding of c-di-GMP signaling and specificity in *M. xanthus*, we used RNA-seq to determine during which stage(s) of the lifecycle the 70 genes encoding c-di-GMP metabolizing enzymes, potential c-di-GMP receptors and known c-di-GMP receptors (from here on “c-di-GMP associated proteins”) are expressed. We found that all these genes are expressed, and with 28 being up-regulated and 10 down-regulated during development. In particular, transcription of the three genes encoding enzymatically active proteins with a stage-specific function were regulated in a stage-specific manner, supporting that temporal regulation of the synthesis of proteins involved in c-di-GMP metabolism contributes to signaling specificity. To inform the RNA-seq analysis, we performed Cappable-seq to identify transcription start sites (TSSs) at a genome-wide scale. These data together with a previously published ChIP-seq analysis to map MrpC binding sites during development (50) revealed nine of the developmentally regulated genes as candidates for being directly regulated by MrpC. In particular, we found that MrpC directly represses *dmxB* and activates *pmxA* expression. Consistently, a Δ*mrpC* mutant has increased accumulation of c-di-GMP and cGAMP.

## Results

### RNA-seq transcriptome profiling reveals pervasive developmental regulation of genes encoding “c-di-GMP associated proteins”

To elucidate whether transcriptional regulation of genes for “c-di-GMP associated proteins” contributes to their stage-specific function, we performed RNA-seq analyses using the wild-type (WT) strain DK1622. To this end, we collected total RNA from non-starved cells (from here on referred to as 0 h of development) and from cells developed for 6, 12, 18 and 24 h under submerged culture conditions. These time points span the entire process of aggregation of cells to form fruiting bodies and the early stages of sporulation. RNA was isolated from two biological replicates. RNA sample preparation, depletion of rRNA, sequencing and data analysis are described in Materials and Methods. Benchmarking of the RNA-seq data using reverse transcription quantitative PCR (RT-qPCR) analyses of the *mrpC* and *fruA* genes that are both transcriptionally up-regulated during development (43, 44, 52, 53) demonstrated that the two genes had the same expression patterns in the two approaches (Fig. S1).

Subsequently, we focused on the 70 genes encoding “c-di-GMP associated proteins. These genes include all those encoding proteins with a GGDEF domain (18), EAL domain (2), HD-GYP domain (6), PilZ domain (24), or MshEN domain (17) as well as CdbA, CdbB and Nla24. All proteins with one of these domains were included because non-enzymatically proteins or proteins that do not bind c-di-GMP can still be involved in regulation of c-di-GMP-dependent processes (11, 58). All 70 genes were expressed with normalized read counts of more than 50 at all five time points (Fig. 1A, Table S1A). A comparison of transcript levels during development to that during growth (0 h), revealed four clusters with distinct expression profiles. One cluster of ten genes including *dmxB, pmxA* and *pkn1* as well as the benchmarking *mrpC* and *fruA* genes were induced more than 4-fold (log_2_FC ≥2, adjusted *p*-value ≤0.05) at one or more time points during development (Fig. 1AB, Table S1B). Pkn1 is a Ser/Thr protein kinase with a C-terminal PilZ domain and is specifically important for development (36, 59); it is not known whether the PilZ domain binds c-di-GMP. These observations are in agreement with previous findings that *dmxB* and *pkn1* transcription is up-regulated during development (40, 59). A second cluster of 18 genes including *tmoK, pixB* and *pilB* were induced more than 2-fold (log_2_FC ≥1, adjusted *p*-value ≤0.05) at one or more time point(s) during development. TmoK is a histidine protein kinase with a C-terminal GGDEF domain and is important for T4P-dependent motility during growth as well as for development; the GGDEF domain does not have DGC activity and does not bind c-di-GMP (37, 40). PilB is the ATPase for T4P extension and contains an N-terminal MshEN domain (17, 60) but it is not known whether it binds c-di-GMP. In the third cluster, ten genes including *dmxA, cdbA* and *cdbB* were down-regulated more than 2-fold (log_2_FC ≥-1, adjusted *p*-value ≤0.05) at one or more time point(s) during development (Fig. 1B). Expression of the remaining 32 genes, including *sgmT, pixA, plpA* and *nla24* were not significantly regulated during development (Fig. 1B, Table S1B). PlpA is a PilZ-domain protein that regulates motility during growth but is not important for development and was reported not to bind c-di-GMP *in vitro* (36, 61). Control experiments using RT-qPCR on the same RNA as for RNA-seq for selected genes (*dmxA, dmxB, pkn1*) reproduced the RNA-seq data (Fig. S1).

**Figure 1.**
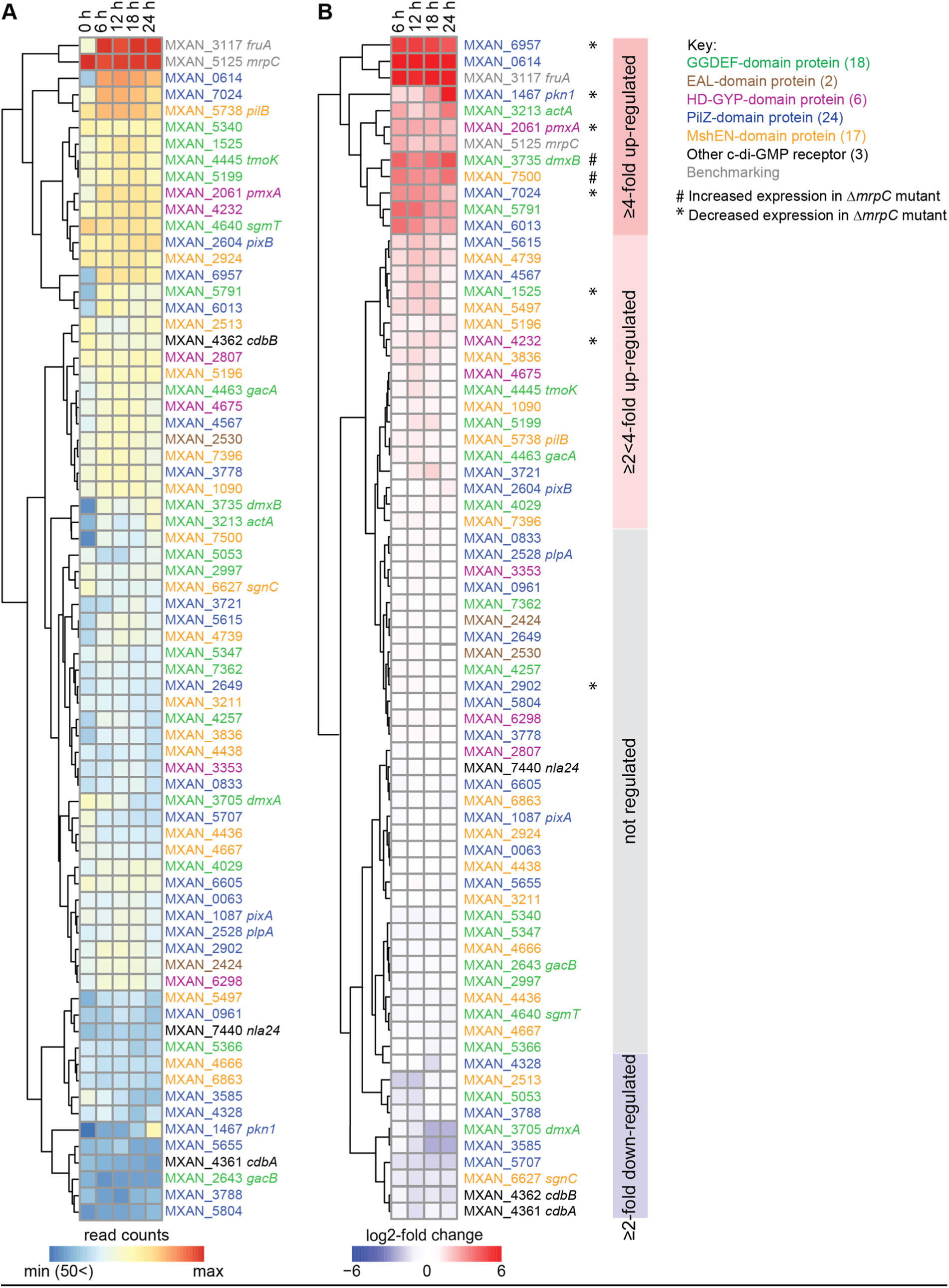
Expression of genes for “c-di-GMP associated proteins”. A. Expression of the genes encoding “c-di-GMP associated proteins”. Heat-map represents normalized read counts at the indicated time points. Genes are colour-coded according to the key on the right. MXAN_2807 is indicated as a protein with an HD-GYP domain; this protein also contains a MshEN domain. B. Relative transcript levels during development for genes encoding “c-di-GMP associated proteins”. Heat-map indicates log2-fold change at 6, 12, 18 or 24 h of development compared to 0 h. Genes marked * or # were expressed at lower and higher levels, respectively in the Δ*mrpC* mutant compared to WT as determined using RT-qPCR (See also Fig. 2 and Fig. S3). Coloured boxed on the right indicate the four clusters with distinct expression profiles.

We conclude that expression of the genes for the enzymatically active proteins (DmxA, DmxB and PmxA) with a stage-specific function correlates with that stage of the lifecycle. Similarly, the gene for the developmentally important Pkn1 protein is up-regulated during development while the gene for the growth-related PixA was constitutively expressed. Similarly, the genes for the three verified c-di-GMP receptors (SgmT, PixB, Nla24) that function during both stages of the lifecycle were expressed constitutively while the genes for the essential proteins CdbA and CdbB were down-regulated during development. Altogether, these observations support that transcriptional regulation of genes encoding proteins that act in a stage-specific manner may contribute to temporally restricting their activity.

### Genome-wide mapping of transcription start sites using Cappable-seq

To further understand transcriptional regulation of genes for “c-di-GMP associated proteins”, we performed genome-wide mapping of transcription start sites (TSSs) with single-nucleotide-resolution using Cappable-seq (62). For this, total RNA was isolated in two biological replicates from growing *M. xanthus* cells (0 h) and from cells developed for 6, 12, 18 and 24 h under the same conditions as for the RNA-seq analysis. RNA samples were enriched for primary transcripts with a triphosphate at the 5’-end and cDNA libraries generated and sequenced (Materials and Methods). The number of reads starting at a certain position was normalized to the total number of reads to obtain a relative read score (RRS) (Materials and Methods) (Table S2). As in (62), TSS with an RRS <1.5 (equivalent to ∼10 reads or less) were discarded from the analysis.

We benchmarked the accuracy of Cappable-seq using the previously mapped TSSs of *fruA* and *mrpC*. For *fruA*, we identified 12 potential TSSs in both biological replicates (Table S3A). The potential TSSs at −235 and −286 relative to the first nucleotide in the start codon (from here on, TSC for translation start codon) were significantly above the threshold and observed at all time points while the remaining 10 were close to the threshold, and generally not observed at all time points. The signal for the TSS at −235 increased during development while the one at −286 did not (Fig. S2A; Table S3A). A TSS at −235 matched the RNA-seq data (Fig. S2A). Importantly, the TSS at −235 matches the previously identified TSS using primer extension on RNA isolated from developing cells (53). For *mrpC*, two potential TSS were identified (Table S3A). The TSS at −58 bp relative to TSC had the highest score, was detected at all time points in both replicates, and increased during development (Fig. S2B; Table S3A). The potential TSS at −21 relative to the TSC was close to the threshold, and detected only at 12 and 24 h. A TSS at −58 matched the RNA-seq data (Fig. S2B; Table S3A). Importantly, a TSS located at −60 bp relative to TSC was identified using primer extension on RNA from developing cells (63). We conclude that Cappable-seq reproduces previously identified TSS of *fruA* and *mrpC* with good accuracy and also identified alternative potential TSSs. These alternative TTSs are likely explained by the higher sensitivity of Cappable-seq compared to primer extension (62). Further work is needed to verify whether they represent genuine TSSs.

### MrpC regulates expression of several genes for “c-di-GMP associated proteins” during development

Having validated the Cappable-seq approach, we aimed to identify the transcriptional units encoding “c-di-GMP associated proteins”. For this, we defined genes likely to be in an operon as those transcribed from the same strand, and with an intergenic distance between stop and start codon of flanking genes ≤50 bp (Table S3C). By combining these data with Cappable-seq data, most genes encoding “c-di-GMP associated proteins” could be divided in four categories: Genes likely not part of an operon (32), likely first gene in an operon (11), likely internal gene in operon (4), and likely internal gene in operon and with an internal promoter (12); for four predicted operons and seven genes predicted not to be in an operon, no TSSs were detected (Table S3BC).

During these analyses, we noticed that several TSSs associated with genes/operons for “c-di-GMP associated proteins” were close to binding site(s) for MrpC as mapped at a genome-wide scale using ChIP-seq on cells developed for 18 h (50). That analysis identified >1500 MrpC binding sites on the *M. xanthus* genome, many of which map to promoter regions of developmentally regulated genes. To identify genes/operons for “c-di-GMP associated proteins” that could potentially be directly regulated by MrpC, we used two criteria. Firstly, we used the criterion of Robinson *et al*. (50) who identified promoter regions with an MrpC binding site as those in which the MrpC ChIP-seq peak was located at a distance of −400 to +100 bp from a TSC. Secondly, based on published experimental data on MrpC binding to the *fruA* and *mrpC* promoters (44, 45, 64), we included the criterion that an MrpC ChIP-seq peak should be located within a distance of 200 bp from a TSS (Fig. S2AB). Based on these criteria, we identified 18 operons/genes for “c-di-GMP associated proteins” that could potentially be regulated by MrpC (Table 3BC). Using RT-qPCR, we found that two (*dmxB* and MXAN_7500) and seven (MXAN_1525, *pmxA*, MXAN_4232, *pkn1*, MXAN_2902, MXAN_6957, MXAN_7024) of these 18 genes were expressed at higher and lower levels, respectively in the Δ*mrpC* mutant compared to WT while nine genes displayed similar expression patterns in the two strains (Fig. 2; Fig. S3). The observation that nine of the candidate genes were not expressed in an MrpC-dependent manner under the conditions tested are in agreement with the possibility that the relevant MrpC ChIP-seq peaks may represent false positives as discussed by Robinson *et al*. We note that the expression of all tested genes in the WT as measured by RT-qPCR matches the expression patterns obtained using RNA-seq (Fig. 1B). The nine differentially expressed genes include six of the most highly developmentally up-regulated genes for “c-di-GMP associated proteins” (Fig. 1B). These results support that MrpC is a negative regulator of *dmxB* and MXAN_7500 expression and a positive regulator of MXAN_1525, *pmxA*, MXAN_4232, *pkn1*, MXAN_2902, MXAN_6957 and MXAN_7024 expression. From here on, we focused on MrpC regulation of *dmxB* and *pmxA*, which encode enzymatically active proteins that are specifically important for development.

**Figure 2.**
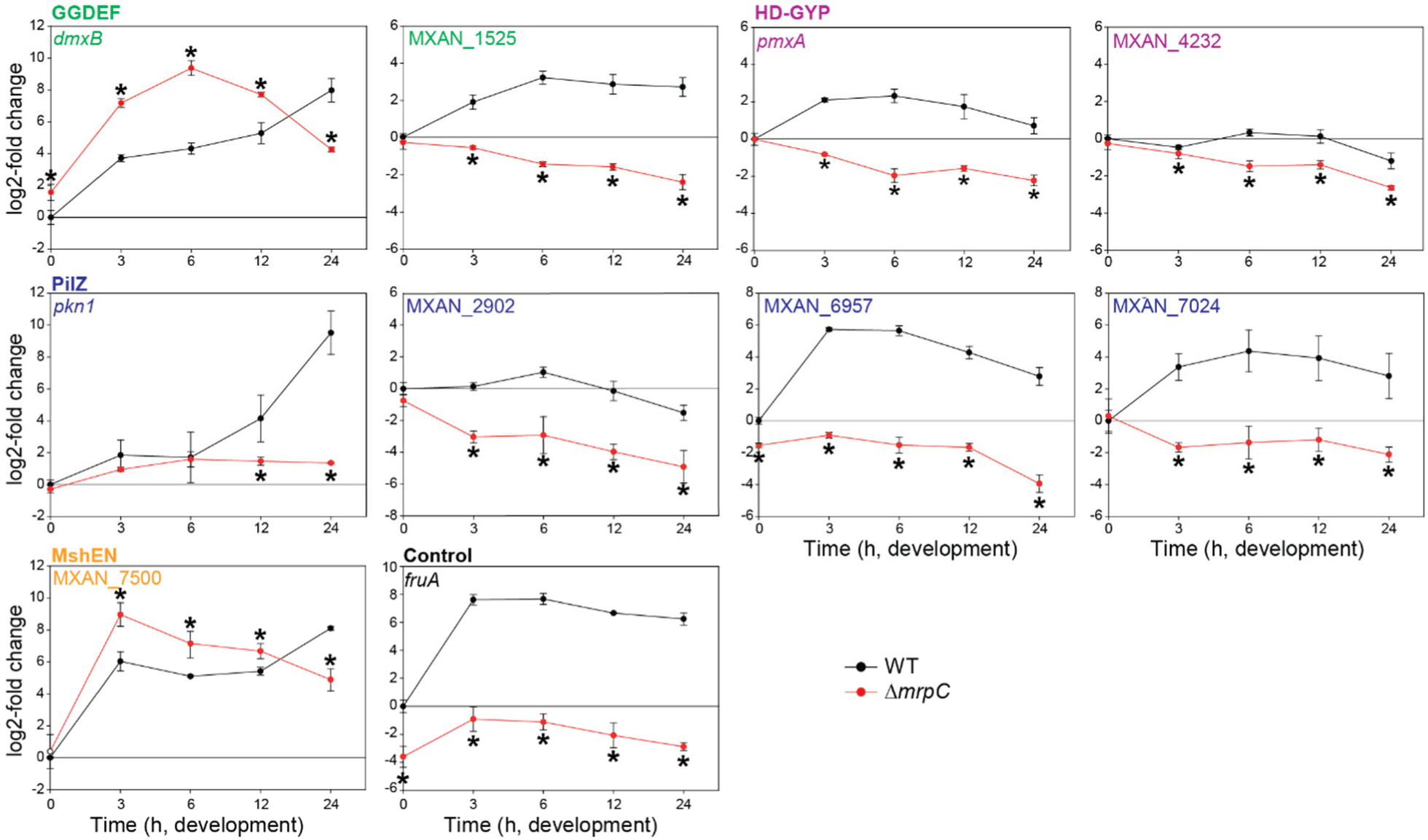
Regulation of expression of genes encoding “c-di-GMP associated proteins” by MrpC. Total RNA was isolated from cells developed in MC7 submerged cultures at the indicated time points from WT (black) and the Δ*mrpC* mutant (red). Transcript levels are shown as mean ± standard deviation (SD) from two biological replicates, each with two technical replicate, relative to WT at 0 h. *, *P*-value <0.05 in Student’s t-test in which samples from the Δ*mrpC* mutant were compared to the samples from WT at the same time point. *fruA* served as a positive control. Based on protein sequence analysis, MXAN_1525 and MXAN_4232 are predicted to have DGC and PDE activity, respectively; however, neither a ΔMXAN_1525 nor a ΔMXAN_4232 mutant has defects during growth or development (37, 40). *pkn1*, MXAN_2902, MXAN_6957 and MXAN_7024 are PilZ-domain proteins; however, none contain the conserved motifs for c-di-GMP binding (27, 36). Except for Pkn1, lack of any of these four proteins does not cause defects during growth or development (36, 59). MXAN_7500 is a MshEN-domain protein with the sequence motifs for c-di-GMP binding (17); however, it is known not whether this protein binds c-di-GMP or whether it is important during growth and development.

### MrpC negatively regulates *dmxB* expression and DmxB accumulation

Based on our criteria as well as RNA-seq, *dmxB* forms a two-gene operon with the downstream gene MXAN_3734 (Fig. 3A; Fig. S4; Table S3BC). We identified seven potential TSSs upstream of *dmxB* in both replicates (Table S3BC). Among these, we focused on four with high scores in both replicates at several time points (Fig. 3B), while the remaining three had low scores and each appeared at only one time point (Table S3BC). The TSS at −297 relative to TSC was detected with the highest score at all time points and increased as development progressed (Fig. 3B, Table S3BC). The TSS at −213 was the second highest scoring and sharply increased at 18 h. The TSSs at −171 and −135 did not significantly change in score over time. A comparison of Cappable-seq and RNA-seq data supports that TSSs at −297 and −213 are genuine TSSs (Fig. 3B; Fig. S4). These data support that *dmxB* is transcribed from multiple promoters, and those with TSSs at −297 and −213 are developmentally regulated.

**Figure 3.**
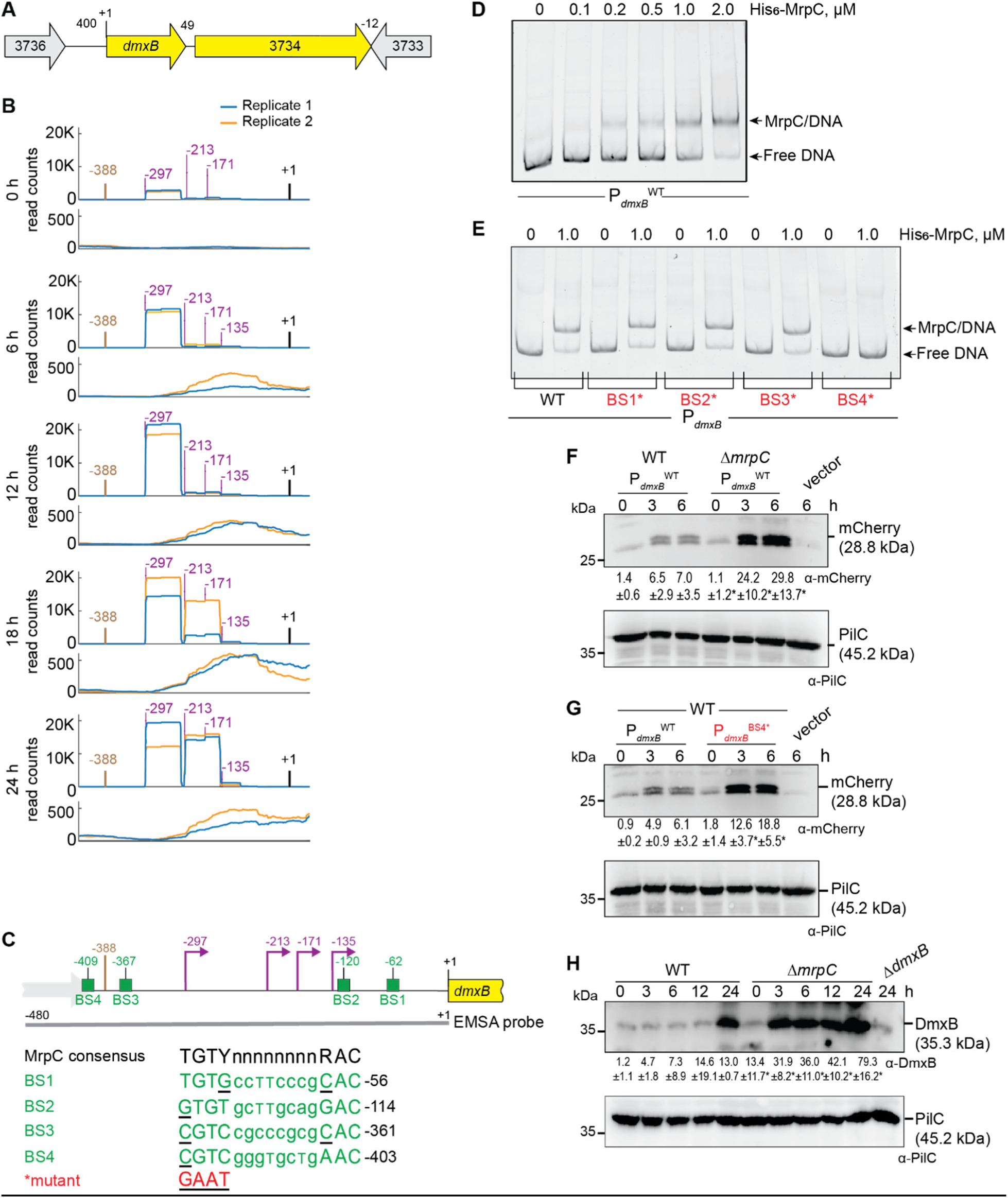
MrpC negatively regulates expression of *dmxB*. A. Schematic of *dmxB* locus. Direction of transcription is indicated by the arrows. +1 indicates TSC of *dmxB*. Numbers above indicate distance between start and stop codons of flanking genes. MXAN_3734 encodes a response regulator that is not important for development (40). B. Visualization of RNA-seq (lower panels) and Cappable-seq (upper panels) data at different time points. For each time point, mapped read counts for both biological replicates are shown in blue and orange. Data from RNA-seq and Cappable-seq are from different samples. +1 indicates the *dmxB* TSC. TSSs as mapped by Cappable-seq are indicated in purple relative to the TSC of *dmxB*. The center of the MrpC ChIP-seq peak is in brown. C. Feature map of *dmxB* promoter region. +1 and colour code is as in B. Green boxes labelled BS1-4 indicate potential MrpC binding sites based on the consensus sequence as defined by (50); sequences of BS1-4 are shown below and in which underlining indicate a mismatch. Red indicates the sequence used to generate the mutant binding sites. D, E. MrpC binds to the *dmxB* promoter region using BS4. The indicated Hex-labelled probes were mixed with the indicated concentrations of His_6_-MrpC EMSA and analyzed by EMSA. F. MrpC represses *dmxB* promoter(s). Total cell lysates from the indicated strains expressing *mcherry* from P_*dmxB*_^WT^ were harvested from cells developed in MC7 submerged cultures at the indicated time points. 10 μg of protein were loaded per lane and samples separated by SDS-PAGE. Upper and lower blots were probed with α-mCherry and α-PilC antibodies, respectively. PilC blot served as loading control. Numbers below upper panel indicate in the accumulation of mCherry relative to PilC as mean ± SD as measured in three biological replicates. *, *P*-value <0.05 in Student’s t-test in which samples from the Δ*mrpC* mutant were compared to samples from WT at the same time point. Vector with *mCherry* but without the *dmxB* promoter served as a negative control (vector). mCherry separates into two bands; the reason for this is not known. G. BS4 is important for MrpC-dependent repression of *dmxB* promoter(s). Total cell lysates from the indicated WT strains expressing mCherry from the two indicated promoters were prepared and analyzed as in F. H. DmxB accumulates at increased levels in the Δ*mrpC* mutant. Total cell lysates of the indicated strains were harvested from cells developed in MC7 submerged conditions at indicated time points and analyzed as in F.

The *dmxB* promoter region contains an MrpC ChIP-seq peak centered at −388 bp relative to TSC (Fig. 3BC; Table S3BC). To test whether MrpC directly binds to the upstream region of *dmxB*, we performed an electrophoretic mobility shift assay (EMSA) using a PCR-amplified 480 bp Hexachloro-fluorescein (Hex)-labeled PCR product that extends from 92 bp upstream of the ChIP-seq peak coordinate to the *dmxB* TSC (Fig. 3C). Titrating purified His_6_-MrpC (Fig. S5) against the Hex-labelled probe resulted in the formation of one well-defined shifted band consistent with one binding site for MrpC in the *dmxB* promoter region (Fig. 3D).

We identified four potential MrpC binding sites (BS1-4) in the *dmxB* promoter region using the consensus sequence defined by (50) (Fig. 3C). We prepared four Hex-labelled *dmxB* promoter fragments each containing substitutions of conserved bp in one of the four potential MrpC binding sites as described (44). In EMSA experiments, the fragments with substitutions in BS1, BS2 or BS3 bound MrpC as the WT fragment (Fig. 3E). By contrast, the fragment with a mutated BS4 did not bind MrpC (Fig. 3E). Based on these data, we suggest that *dmxB* promoter contains one binding site, i.e. BS4, for MrpC centered at −409 and, thus, close to the MrpC ChIP-seq peak centered at −388 bp (Fig. 3C).

To test the impact of MrpC and its binding to BS4 on *dmxB* promoter activity *in vivo*, we constructed fluorescent reporters in which the WT *dmxB* promoter fragment (P_*dmxB*_^WT^) used in the EMSA experiments or the same fragment with a mutated BS4 (P_*dmxB*_^BS4*^) were fused to the start codon of *mCherry* and ectopically expressed from the Mx8 *attB* site. As a negative control, *mCherry* without the *dmxB* promoter was fused to *mCherry*. In agreement with the RT-qPCR data (Fig. 2), mCherry expressed from P_*dmxB*_^WT^ accumulated at significantly higher levels in the Δ*mrpC* mutant compared to WT at all tested time points (Fig. 3F). Importantly, the activity of P_*dmxB*_^BS4*^ was significantly higher than that of P_*dmxB*_^WT^ in WT (Fig. 3G). We conclude that MrpC binds to BS4 to repress *dmxB* expression.

Finally, we observed that DmxB was detected at low levels at 0 h in WT and its accumulation increased during development (Fig. 3H) as previously observed (40). Importantly, DmxB accumulated at significantly higher levels at all time points in the Δ*mrpC* mutant compared to WT (Fig. 3H) consistent with MrpC acting as a repressor of *dmxB* transcription.

### MrpC positively regulates *pmxA* expression and PmxA accumulation

Based on our criteria, *pmxA* is the last gene of a three gene operon (Fig. 4A). Based on Cappable-seq, there is one TSS at −63 relative to the TSC of MXAN_2063, and three TSSs immediately upstream of *pmxA* (Fig. 4B; Table S3BC). RT-PCR analysis on RNA isolated from WT at 0 and 6 h of development support that MXAN_2063-MXAN_2062-*pmxA* is transcribed as an operon at both time points (Fig. S6A). The three genes were barely expressed at 0 h; at later time points, MXAN_2063 and MXAN_2062 expression remained low while *pmxA* expression increased (Fig. 4B; Fig. S6B). Accordingly, the score for the single TSS upstream of MXAN_2063 remained low (Fig. 4B; Table S3C). The TSSs upstream of *pmxA* had scores close to the threshold (Table S3BC). Therefore, we analyzed each biological replicate separately (Fig. 4B, right panels; Table S3BC). A TSS at −226 relative to the TSC of *pmxA* was detected at all time points and was not developmentally regulated while a TSS at −131 was detected at 6 h and later suggesting developmental up-regulation. A TSS at −53 was detected only at 24 h. We conclude that the MXAN_2063-MXAN-2062-*pmxA* operon is transcribed from a promoter upstream of MXAN_2063 during growth and development; in addition, *pmxA* is transcribed from internal promoters, two of which are developmentally regulated.

**Figure 4.**
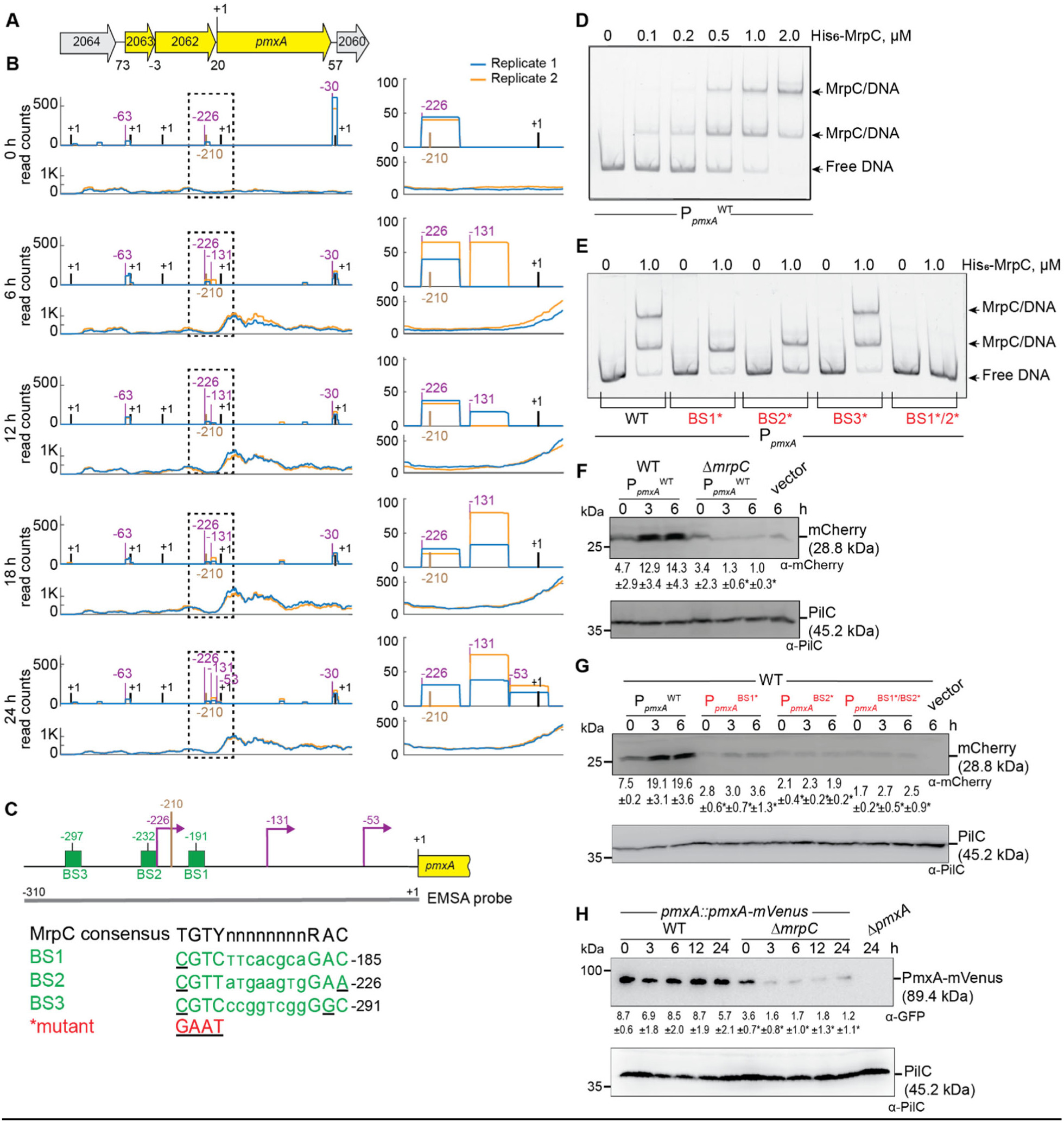
MrpC positively regulates expression of *pmxA*. A. Schematic of *pmxA* locus. Direction of transcription is indicated by the arrows. +1 indicates TSC of *pmxA*. Numbers above indicate distance between start and stop codons of flanking genes. MXAN_2063 encodes a FecR domain-containing protein with a lipoprotein signal peptide and MXAN_2062 encodes a protein with a type I signal peptide, an N-terminal LysM domain and a C-terminal extracellular fibronectin type III domain. The function of these two proteins is not known. B. Visualization of RNA-seq (lower panels) and Cappable-seq (upper panels) data at different time points for genes at *pmxA* locus. For each time point, mapped read counts for both biological replicates are shown in blue and orange. The data from RNA-seq and Cappable-seq were obtained from different samples. Left panels, +1 indicates TSCs of MXAN_2064-_2060; right panels, zoom of region indicated in the hatched box in left panels immediately upstream of *pmxA* and where +1 indicates the TSC of *pmxA*. In both sets of panels, TSSs as mapped by Cappable-seq are indicated in purple relative to the nearest TSC. The center of the MrpC ChIP-seq peak is in brown. C. Feature map of *pmxA* promoter region. +1 and colour code is as in B. Green boxes labelled BS1-3 indicate potential MrpC binding sites based on the consensus sequence as defined by (50); sequences of BS1-3 are shown below and in which underlining indicate a mismatch. Red indicates the sequence used to generate the mutant binding sites. D, E. MrpC binds to the *pmxA* promoter region using BS1 and BS2. The indicated Hex-labelled probes were mixed with the indicated concentrations of His_6_-MrpC EMSA and analyzed by EMSA. F. MrpC activates *pmxA* promoter(s). Total cell lysates from the indicated strains expressing *mcherry* from P_*pmxA*_^WT^ were harvested from cells developed in MC7 submerged cultures at the indicated time points and then analyzed as in Fig. 3F. G. BS1 and BS2 are important for MrpC-dependent activation of the *pmxA* promoter(s). Total cell lysates from the indicated WT strains expressing mCherry from the indicated promoters were prepared and analyzed as in Fig. 3F. H. PmxA accumulates at reduced levels in the Δ*mrpC* mutant. Total cell lysates of the indicated strains were harvested from cells developed in MC7 submerged conditions at indicated time points and analyzed as in Fig. 3F.

We identified a single MrpC ChIP-seq peak centered at −210 upstream of the *pmxA* TSC and none upstream of MXAN_2063 suggesting that MrpC is involved in activation of the internal promoter(s) during development (Fig. 4BC). In EMSA experiments with a 310 bp Hex-labeled probe (Fig. 4C), 0.1 μM His_6_-MrpC gave rise to a single well-defined shifted band, and at 0.5-2.0 μM His_6_-MrpC, an additional well-defined shifted band was evident (Fig. 4D). We identified three potential MrpC binding sites (BS1-3) upstream of *pmxA* (Fig. 4C), mutated them separately, and tested His_6_-MrpC binding to the mutated promoters. The P_*pmxA*_^WT^ fragment gave rise to two shifted bands at 1.0 μM His_6_-MrpC, while the fragments containing substitutions in BS1 or BS2 generated only one shifted band, the fragment with substitutions in BS3 behaved as P_*pmxA*_^WT^, and a fragment with both BS1 and BS2 mutated did not bind MrpC at 1.0 μM (Fig. 4C, 4E). We conclude that MrpC binds to the internal *pmxA* promoter region at two sites, BS1 and BS2, centered at −191 and −232 relative to the TSC of *pmxA* (Fig. 4C).

The importance of MrpC and its binding to BS1 and BS2 on *pmxA* promoter activity *in vivo* was tested as described for P_*dmxB*_ using the same fragments as in the EMSA experiments. mCherry expressed from P_*pmxA*_^WT^ was detected in immuno-blots of WT at 0, 3 and 6 h, and at significantly reduced levels in the Δ*mrpC* mutant at 3 and 6 h (Fig. 4F) in agreement with the RT-qPCR experiments (Fig. 2). Importantly, the activity of P_*pmxA*_^BS1*^, P_*pmxA*_^BS2*^ and P_*pmxA*_^BS1*/BS2*^ was significantly lower than that of P_*pmxA*_^WT^ in WT (Fig. 4G).

To determine PmxA levels during development, we used an active PmxA-mVenus fusion (Fig. S6C) expressed from the native site. Surprisingly, the level of PmxA-mVenus did not increase significantly during development in WT (Fig. 4H) despite transcription being up-regulated ∼4-fold during development (Fig. 1B, 2B and Fig. 4F). Importantly, the level of PmxA-mVenus in the Δ*mrpC* mutant was significantly reduced compared to WT at all time points (Fig. 4H). Altogether these observations support that *pmxA* is transcribed from a promoter upstream of MXAN_2063 as well as from internal promoter(s), which are activated by MrpC by binding to BS1 and BS2.

### MrpC curbs accumulation of c-di-GMP and 3’, 3’ cGAMP during development

Next, we investigated the functional consequences of altered accumulation of DmxB and PmxA with respect to cyclic di-nucleotides in the Δ*mrpC* mutant. As described (40), the c-di-GMP level increased significantly during development in a DmxB-dependent manner in the WT (Fig. 5A). In agreement with the accumulation profile of DmxB, the c-di-GMP level was slightly but significantly higher in the Δ*mrpC* mutant than in the WT at 0 h, and significantly higher during development in the Δ*mrpC* mutant, and the extra c-di-GMP was dependent on DmxB (Fig. 5A).

**Figure 5.**
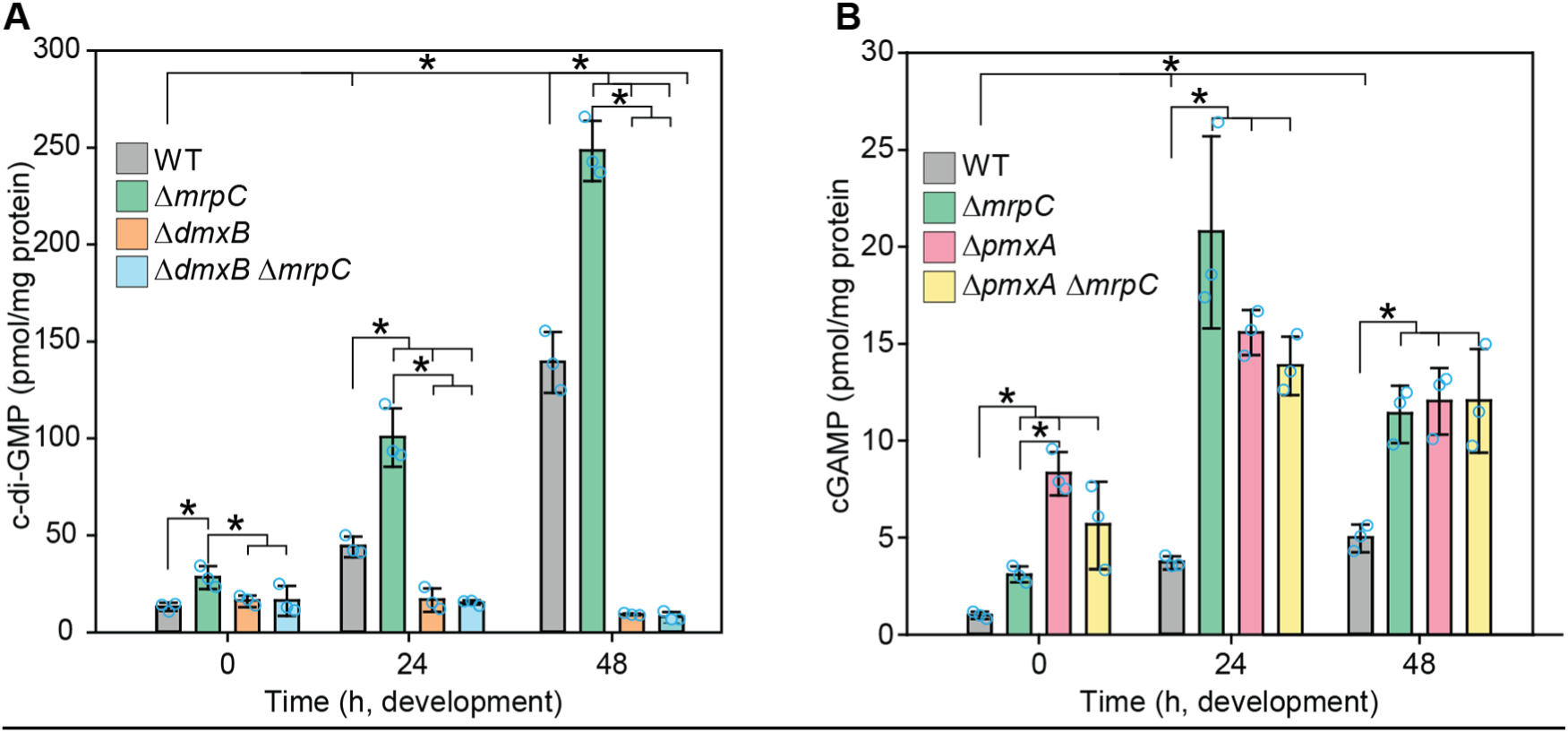
c-di-GMP and cGAMP accumulation in WT and *ΔmrpC* mutant during development. A, B. c-di-GMP and cGAMP levels during growth and development. Cells were harvested at the indicated time points of development, and nucleotide levels and protein concentrations determined. Levels are shown as mean±SD calculated from three biological replicates. Individual data points are in light blue. *, *P*-value <0.05 in Student’s t-test.

Because recent studies revealed that PmxA activity against c-di-GMP is significantly lower than towards cGAMP (54), we measured c-di-GMP as well as cGAMP levels in WT and the Δ*pmxA* and Δ*mrpC* mutants. As previously shown (40), the c-di-GMP level in the WT and the Δ*pmxA* mutant were similar (Fig. S7). The cGAMP level increased significantly during development in WT (Fig. 5B). Importantly, in the Δ*pmxA* mutant, the cGAMP level was significantly higher than in WT during growth (0 h) as well as development, consistent with the accumulation profile of PmxA-mVenus and PmxA having PDE activity against cGAMP *in vivo* (Fig. 4H). The cGAMP level in the Δ*mrpC* mutant was significantly higher than in WT at all time points and, except at 0 h, largely similar to that in the Δ*pmxA* mutant. Finally, the Δ*pmxA*Δ*mrpC* mutant accumulated cGAMP similarly to the Δ*pmxA* mutant documenting that the increased cGAMP level in the Δ*mrpC* mutant depends on PmxA.

We conclude that MrpC by regulating the expression of *dmxB* and *pmxA* controls the cellular pools of c-di-GMP and cGAMP.

### Aggregated and non-aggregated cells accumulate MrpC, DmxB, PmxA-mVenus as well as c-di-GMP or cGAMP at similar levels

In the DZ2 WT strain, MrpC expression and accumulation are higher in aggregated cells, i.e. cells that differentiate to spores within fruiting bodies, compared to non-aggregated cells, i.e. cells that differentiate to peripheral rods (33, 44) raising the possibility that c-di-GMP and/or cGAMP might also accumulate at different levels in these cell types. To this end, we developed DK1622 WT cells under submerged conditions and then separated aggregated and non-aggregated cells at 24 and 48 h of development and determined MrpC, DmxB, PmxA-mVenus, c-di-GMP and cGAMP levels in the two cell types. As a control for proper cell separation, we used accumulation of Protein C, which accumulates in aggregated cells and at a much-reduced level in non-aggregated cells (65). In WT as well as in WT producing PmxA-mVenus, cells were properly separated based on the level of Protein C (Fig. 6A). Surprisingly, at both time points, MrpC accumulated at similar levels in the two cell types (Fig. 6A). Consistently DmxB and PmxA-mVenus accumulated at similar levels in the two cell types (Fig. 6A) and c-di-GMP (Fig. 6B) as well as cGAMP (Fig. 6C) levels were similar in the two cell types at both time points. These observations support that MrpC, DmxB, PmxA, c-di-GMP and cGAMP are not involved in cell fate determination during development in DK1622 WT.

**Figure 6.**
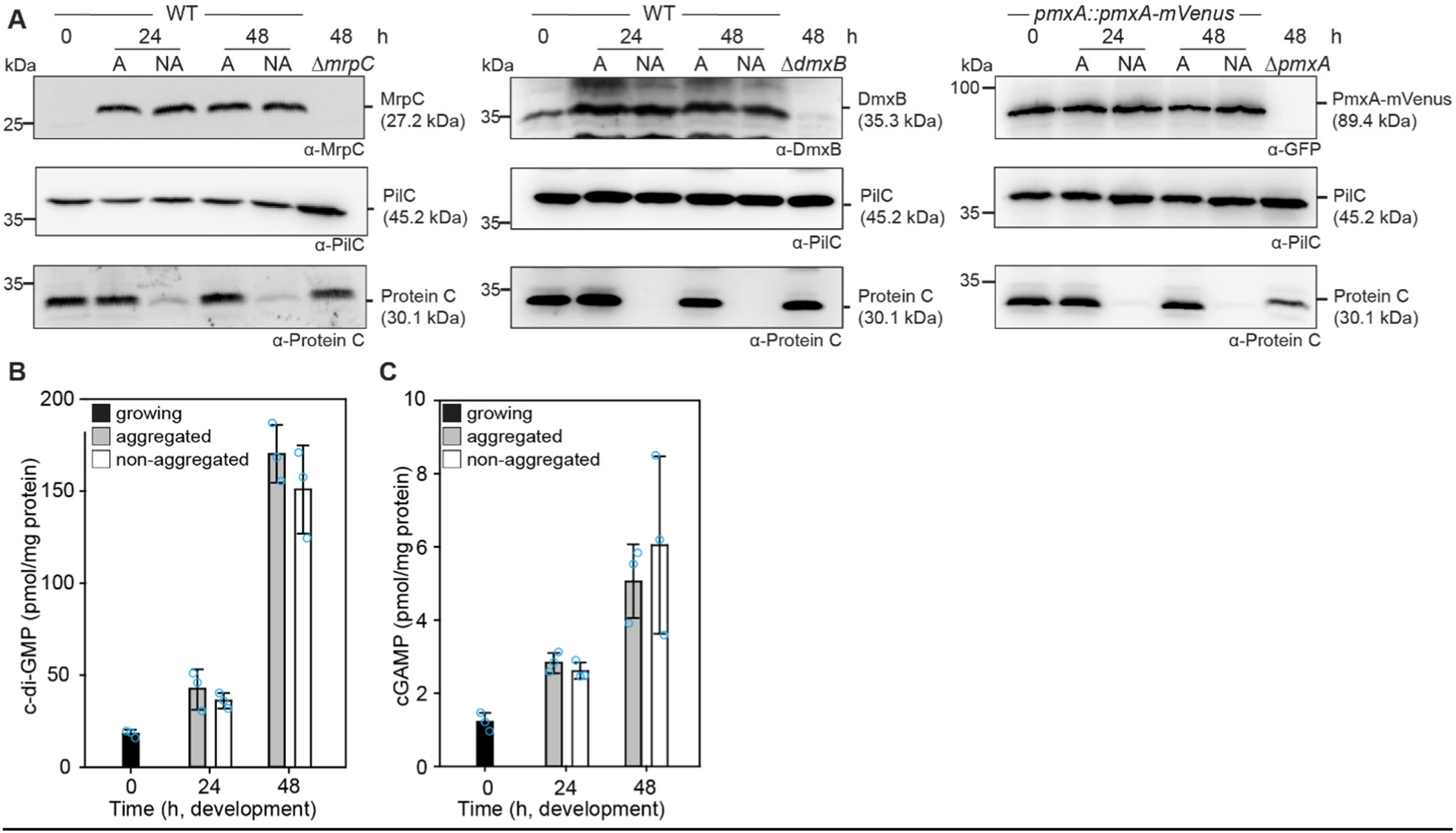
MrpC, DmxA, PmxA-mVenus, c-di-GMP and cGAMP accumulation in aggregated and non-aggregated cells. A. MrpC, DmxB and PmxA-mVenus accumulate at the same levels in aggregated and non-aggregated cells. Cells were harvested at the indicated time points of development and separated into the two cell fractions. 10 μg of protein was loaded per lane and samples separated by SDS-PAGE. Upper blots were probed with α-MrpC, α-DmxB or α-GFP, middle blots with α-PilC, and lower blots with α-Protein C antibodies. The PilC blots served as loading controls and the Protein C blots as cell separation controls. B, C. c-di-GMP (B) and cGAMP (C) accumulate at the same levels in aggregated and non-aggregated cells of WT. Samples were generated as in A. Levels are shown as mean±SD calculated from three biological replicates. Individual data points are in light blue. *, *P*-value <0.05 in Student’s t-test.

## Discussion

Here, we present a comprehensive analysis of the expression of genes encoding “c-di-GMP associated proteins” in *M. xanthus*. This analysis was motivated by previous observations that lack of several of these proteins cause defects during only one of the stages of the biphasic life cycle while others cause defects during both stages. Using RNA-seq, we found that all these genes were expressed during the lifecycle. More importantly, expression of 28 genes encoding “c-di-GMP associated proteins” was up-regulated, 10 down-regulated, and 32 did not change expression during development. By combining Cappable-seq with data from previously published ChIP-Seq analyses of the CRP-like transcription factor MrpC (50), we identified nine genes for “c-di-GMP associated proteins” that are regulated (directly or indirectly) by MrpC. Among these, detailed analyses revealed that (1) MrpC binds to and represses the promoter(s) of *dmxB*, which encodes the DGC DmxB that is essential for development and responsible for the dramatic increase in c-di-GMP during development; and (2) MrpC binds to and activates internal promoter(s) in the MXAN_2063-MXAN_2062_*pmxA* operon to promote transcription of *pmxA*, which encodes a PDE that is essential for development. Thereby, MrpC regulates the cellular pools of c-di-GMP and cGAMP. Altogether, our findings support that differential expression of genes for “c-di-GMP associated proteins” contribute to their stage-specific function. Moreover, we conclude that MrpC is important for the temporal regulation of genes for c-di-GMP synthesis and cGAMP degradation, and a key regulator of cyclic nucleotide metabolism in *M. xanthus*.

Expression of *dmxB* and DmxB accumulation are up-regulated during development (37, 40). Consistently, lack of DmxB DGC activity only causes developmental defects. We found that *dmxB* is likely expressed from four promoters, two of which are developmentally up-regulated and two constitutively expressed at low levels (Fig. 7). MrpC is not important for up-regulation of *dmxB* transcription during development; rather MrpC represses transcription of *dmxB* during growth and development. Based on EMSA analyses, MrpC binds to a single site (BS4) centered at −409 relative to the TSC to accomplish this function. The MrpC binding site is located 112, 196, 238 and 274 bp upstream from the four TSSs (Fig. 7); however, from our current analyses, we do not know which promoter(s) is repressed by MrpC. The distance between the MrpC binding sites and the four TSSs strongly argues that MrpC does not directly block binding of the RNA polymerase. Recently, McLaughlin *et al*. (44) elegantly demonstrated that MrpC functions as a negative autoregulator of the *mrpC* promoter by outcompeting binding of the MrpB transcriptional activator, which is an enhancer binding protein. We speculate that MrpC may function by a similar mechanism in *dmxB* expression. However, the activator of *dmxB* developmental expression remains to be identified. The MrpC-dependent repression of *dmxB* expression curbs DmxB synthesis and, consequently, c-di-GMP accumulation slightly during growth and more significantly during development. We previously showed that an increase in the global pool of c-di-GMP is essential for development; however, further increasing this level does not interfere with development (40) arguing that the increased c-di-GMP pool in the Δ*mrpC* mutant may not significantly contribute to the developmental defects in this mutant. Rather we suggest that the importance of the negative regulation of *dmxB* expression by MrpC lies in avoiding futile synthesis of DmxB and c-di-GMP.

**Figure 7.**
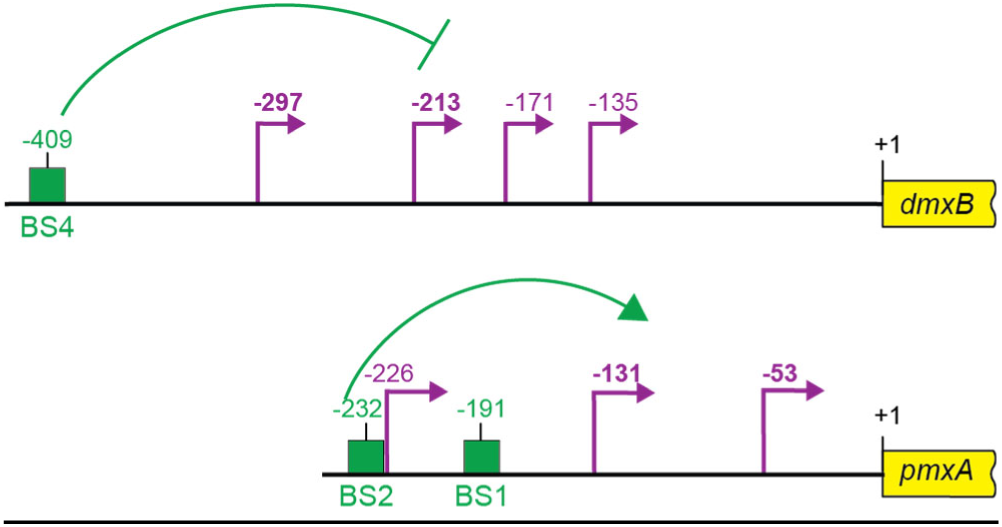
Schematic of *dmxB* and *pmxA* promoter regions. +1 indicate TSC of *dmxB* or *pmxA*; potential TSSs are indicated in purple with developmentally regulated TSSs in bold; green boxes indicate verified MrpC binding sites named as in Fig. 3C and 4C. All coordinates are relative to the TSC (+1).

Lack of PmxA only causes developmental defects. Consistently, expression of *pmxA* is up-regulated during development. *pmxA* is part of a three gene operon, which is expressed a low levels during growth and development. In addition, *pmxA* is expressed from three internal promoters, two of which are developmentally up-regulated (Fig. 7). Our data suggest that the developmental up-regulation of *pmxA* expression derives from the internal promoters. MrpC is essential for up-regulation of *pmxA* transcription; and, based on EMSA analyses, MrpC binds to two sites (BS1 and BS2) centered at −191 and −232 relative to the TSC. Because BS1 only has one mis-match compared to the consensus MrpC binding site while BS2 has two (Fig. 4C), we suggest that MrpC binds BS1 with a higher affinity than BS2. From our current analyses, we do not know which of the internal promoters are activated by MrpC. However, based on a comparison to CRP-activated promoters in *Escherichia coli* (66) and the distance between the MrpC binding sites and the TSSs, we speculate that the promoter with a TSS at −131 relative to the TSC could be activated by MrpC. In the case of the promoter with a TSS at −53, the distance to the MrpC binding sites makes it less likely that this promoter is directly activated by MrpC; however, we notice that CRP in *Escherichia coli* can act as a as a structural element from long distances together with an additional transcriptional activator as in the case of the *malK* promoter (67). It is also a possibility that the promoter with a TSS at −53 is activated by MrpC together with FruA as described for several developmentally regulated promoters (46-51). While transcription of *pmxA* is up-regulated during development in WT, the level of PmxA accumulation (as measured using an active PmxA-mVenus fusion) does not change significantly. By contrast, in the Δ*mrpC* mutant, *pmxA* transcription is not up-regulated and PmxA accumulation is strongly decreased. These observations indicate that PmxA accumulation is not only regulated at the transcriptional level but also at the translational and/or post-translational level. PmxA is essential for development arguing that the reduced *pmxA* expression and PmxA accumulation in the Δ*mrpC* mutant contributes to the developmental defects in this mutant. However, the developmental defects of the Δ*mrpC* mutant are more severe than in the case of the Δ*pmxA* mutant (43, 68) supporting that reduced PmxA accumulation alone does not explain the developmental defects in the Δ*mrpC* mutant.

PmxA is a PDE with higher activity towards cGAMP than c-di-GMP (40, 54). Accordingly, the cellular pool of c-di-GMP is unaltered in a Δ*pmxA* mutant compared to WT. We found that the level of cGAMP increased during development of WT; importantly, the cGAMP level was significantly higher in the Δ*pmxA* mutant compared to WT. Similarly, we found that the cGAMP pool is highly increased in the Δ*mrpC* mutant. These data for the first time show that cGAMP accumulates in *M. xanthus in vivo* and also provide evidence that PmxA is directly involved in its degradation *in vivo*. We speculate that a low concentration of cGAMP maintained by PmxA might be important for development. In *M. xanthus*, GacA and GacB both belong to the Hypr subfamily of GGDEF domain proteins that synthesize cGAMP rather than c-di-GMP *in vitro* (69). Lack of GacA or GacB does not cause evident phenotypes during growth and development (37, 40) but the cGAMP level in these mutants is not known. Based on the RNA-seq data, *gacA* is up-regulated two-fold during development while *gacB* is constitutively expressed (Fig. 1B) and none of these two genes appear to be regulated by MrpC. In future experiments, it will interesting to analyze development and the cGAMP level in a Δ*gacA* Δ*gacB* double mutant to determine whether cGAMP is important for development.

In addition to *dmxB* and *pmxA*, MrpC positively or negatively regulates expression of seven genes for “c-di-GMP signaling proteins” during development (Fig. 1B, Fig. 2). Among these, only the gene for Pkn1, which is up-regulated in an MrpC-dependent manner during development, has been shown to be important for development and none for growth (36, 59) suggesting that lack of Pkn1 may also contribute to the developmental defects in the Δ*mrpC* mutant. Interestingly, we found that some of the MrpC-regulated genes are also differentially expressed during growth. Along these lines, DmxB and PmxA accumulation was increased and decreased, respectively and the levels of c-di-GMP and cGAMP increased during growth in the Δ*mrpC* mutant. The significance of these observations is not clear because lack of MrpC was reported to only cause developmental defects (43). Nevertheless, they indicate that MrpC accumulates during growth but has its primary function in development.

The DGC DmxA is only important during growth (37, 40) and its gene is transcriptionally down-regulated during development. Based on the mapped MrpC ChIP-seq peaks, this down-regulation is independent of MrpC. The reciprocal regulation of *dmxA* and *dmxB* together with the up-regulation of *pmxA*, support a model whereby the signaling specificity of enzymatically active DGCs and PDE with discrete functions during growth and development relies on their temporally regulated synthesis. By contrast, no clear picture emerges for the experimentally verified c-di-GMP receptors regarding transcription of the involved genes: The genes for Nla24, SgmT and PixB that all function during growth and development, are constitutively expressed (*nla24* and *sgmT*) or up-regulated (*pixB*); the gene for PixA, which functions during growth, is constitutively expressed. Clearly, more work is needed to understand how these receptors are regulated and their function restricted to certain stages of the lifecycle.

During development, *M. xanthus* adopts three different cell fates, i.e. peripheral rods, spores or cell lysis. Previous experiments using the WT strain DZ2 demonstrated that MrpC accumulates in aggregated cells that differentiate to spores but at a much-reduced level in non-aggregated cells that differentiate to peripheral rods (33). Because c-di-GMP drives cell fate determination in *Caulobacter crescentus* (70), we speculated that c-di-GMP and/or cGAMP could also play a role in cell fate determination in *M. xanthus*. We found that developing cells of the WT strain DK1622 also segregate into aggregated and non-aggregated cells based on the cell type-specific accumulation of Protein C; however, in this WT strain, MrpC as well as DmxB, PmxA-mVenus, c-di-GMP and cGAMP accumulated at similar levels in the two cell types. These observations argue that MrpC, DmxB, PmxA, c-di-GMP and cGAMP are not involved in cell fate determination during development in DK1622.

## Acknowledgements

We thank Dr. Dobromir Szadkowski for helping with MATLAB scripts, and Dr. Dorota Skotnicka as well as Dr. Anke Treuner-Lange for many helpful discussions. We deeply acknowledge the assistance of the Research Service Centre Metabolomics at the Hannover Medical School in the determination of c-di-GMP and cGAMP levels and the Max Planck Genome Centre Cologne (https://mpgc.mpipz.mpg.de/home) for RNA-seq library construction and sequencing. We acknowledge technical assistance by the Bioinformatics Core Facility at the professorship of Systems Biology at JLU Giessen and the provision of computer resources and general support by the BiGi service center (BMBF grant 031A533) within the de.NBI network. This work was also generously supported by the Deutsche Forschungsgemeinschaft (DFG; German Research Council) within the framework the SPP1879 “Nucleotide second messenger signaling in bacteria” as well as by the Max Planck Society.

## Data Availability

All data supporting this study are available within the article, the Supplementary Information files, or at EBI Arrayexpress (http://www.ebi.ac.uk/arrayexpress; RNA-Seq E-MTAB-11043, Cappable-Seq E-MTAB-11042).

## Code availability

Code for the Cappable-Seq analysis and the Curare version used for the RNA-Seq analysis can be found at Zenodo (www.zenodo.org, ID: 5541852).

## Author contributions

S.K., D.M., A.B. and L.S.-A. conceptualized the study.

S.K., P.B., D.M. and A.G. performed bioinformatics studies.

S.K. performed genetic and molecular microbiology experiments.

S.K. and L.S.-A. wrote the original draft of the manuscript.

All authors reviewed and edited the original manuscript and approved the final version of the manuscript.

AB, AG and L.S.-A. acquired funding and provided supervision.

## Materials and Methods

### Cultivation of *M*. *xanthus* and *E*. *coli*

All *M. xanthus* strains used in this study are derivatives of WT DK1622 (71). In-frame deletions were generated as described (72). All plasmids were verified by sequencing. All strains were confirmed by PCR. *M. xanthus* strains, plasmids and oligonucleotides used are listed in Table 1, Table 2, and Table S4, respectively. *M. xanthus* cells were grown at 32°C in 1% CTT broth (1% Bacto Casitone (Gibco), 10 mM Tris-HCl pH 8.0, 1 mM KPO_4_ pH 7.6, 8 mM MgSO_4_) (73) or on 1% CTT 1.5% agar plates with addition of kanamycin (40 μg ml^-1^) or oxytetracycline (10 μg ml^-1^) if relevant. *E. coli* cells were cultivated in LB (74) or on LB 1.5% agar plates at 37ºC with addition of kanamycin (40 μg ml^-1^) or tetracycline (10 μg ml^-1^) if relevant. All plasmids were propagated in *E. coli* Top10 (Invitrogen™ life technologies) unless otherwise mentioned.

**Table 1.** *M. xanthus* strains used in this study

| Strain | Characteristics | Reference |
| --- | --- | --- |
| DK1622 | Wild-type (WT) | (71) |
| SA5605 | $\Delta dmxB$ | (37) |
| SA3546 | $\Delta pmxA$ | (37) |
| SA6462 | $\Delta mrpC$ | (36) |
| SA8038 | $pmxA::pmxA$-mVenus | This study |
| SA8044 | $\Delta mrpC$, $pmxA::pmxA$-mVenus | This study |
| SA8096 | $attB::pSK65$ (mCherry) | This study |
| SA8098 | $attB::pSK81$ ($P_{pmxA}$-mCherry) | This study |
| SA10108 | $attB::pSK103$ ($P_{pmxA}^{BS1*}$-mCherry) | This study |
| SA10109 | $attB::pSK105$ ($P_{pmxA}^{BS2*}$-mCherry) | This study |
| SA10111 | $attB::pSK111$ ($P_{pmxA}^{BS1*/BS2*}$-mCherry) | This study |
| SA8099 | $attB::pSK101$ ($P_{dmxB}$-mCherry) | This study |
| SA10110 | $attB::pSK112$ ($P_{dmxB}^{BS4*}$-mCherry) | This study |
| SA10133 | $\Delta dmxB \Delta mrpC$ | This study |
| SA10113 | $\Delta mrpC attB::pSK81$ ($P_{pmxA}$-mCherry) | This study |
| SA10105 | $\Delta mrpC attB::pSK101$ ($P_{dmxB}$-mCherry) | This study |
| SA8037 | $\Delta pmxA \Delta mrpC$ | This study |

**Table 2.** Plasmids used in this study

| Plasmid | Description | Reference |
| --- | --- | --- |
| pBJ114 | *galK*, Kan<sup>R</sup> | (86) |
| pSWU30 | *attP*, Tet<sup>R</sup> | (87) |
| pPH158 | pET28a(+), His<sub>6</sub>-*mrpC*, Kan<sup>R</sup> | (33) |
| pSK29 | pBJ114, *pmxA*-mVenus, gene replacement at native site, Kan<sup>R</sup> | This study |
| pSK65 | pSWU30, mCherry, *attB*, Tc<sup>R</sup> | This study |
| pSK81 | pSWU30, P<sub>*pmxA*</sub>-mCherry, *attB*, Tc<sup>R</sup> | This study |
| pSK103 | pSWU30, P<sub>*pmxA*</sub><sup>BS1*</sup>-mCherry, *attB*, Tc<sup>R</sup> | This study |
| pSK105 | pSWU30, P<sub>*pmxA*</sub><sup>BS2*</sup>-mCherry, *attB*, Tc<sup>R</sup> | This study |
| pSK114 | pSWU30, P<sub>*pmxA*</sub><sup>BS3*</sup>-mCherry, *attB*, Tc<sup>R</sup> | This study |
| pSK111 | pSWU30, P<sub>*pmxA*</sub><sup>BS1*/BS2*</sup>-mCherry, *attB*, Tc<sup>R</sup> | This study |
| pSK101 | pSWU30, P<sub>*dmxB*</sub>-mCherry, *attB*, Tc<sup>R</sup> | This study |
| pSK121 | pSWU30, P<sub>*dmxB*</sub><sup>BS1*</sup>-mCherry, *attB*, Tc<sup>R</sup> | This study |
| pSK115 | pSWU30, P<sub>*dmxB*</sub><sup>BS2*</sup>-mCherry, *attB*, Tc<sup>R</sup> | This study |
| pSK109 | pSWU30, P<sub>*dmxB*</sub><sup>BS3*</sup>-mCherry, *attB*, Tc<sup>R</sup> | This study |
| pSK112 | pSWU30, P<sub>*dmxB*</sub><sup>BS4*</sup>-mCherry, *attB*, Tc<sup>R</sup> | This study |

### Development under submerged conditions and cell separation

Exponentially growing *M. xanthus* in CTT were harvested at 5,000 *g* for 5 min and resuspended in MC7 buffer (10 mM MOPS pH 6.8, 1 mM CaCl_2_) to 7×10^9^ cells ml^-1^. 1 ml of concentrated cells was added to 10 ml of MC7 buffer in a polystyrene Petri dish with a diameter of 9.2 cm (Sarstedt). For separation of aggregated and non-aggregated cells during development, cells were developed as described and separated following the procedure of (33). Cells were visualized using a Leica DMi8 inverted microscope with Leica DFC280 camera. To determine sporulation efficiency, cells at 120 h of development were harvested, sonicated for 1 min (30% pulse; 50% amplitude with a UP200St sonifier and microtip; Hielscher) to disperse fruiting bodies and then incubated at 55°C for 2 h. Sporulation efficiency was calculated as the number of sonication- and heat-resistant spores formed after 120 h of development, relative to the WT. Spores were counted in a counting chamber (depth, 0.02mm; Hawksley).

### RNA sequencing

Total RNA from *M. xanthus* cells developed under submerged conditions was extracted from cells using TRI Reagent (Sigma-Aldrich) according to the manufacturer’s protocol. Purified RNA was treated with TURBO DNA-free™ Kit (Invitrogen) according to the manufacturer’s protocol. RNA integrity was analyzed by 1% agarose gel electrophoresis. For all samples rRNA depletion, library preparation and sequencing were performed at the Max-Planck-Genome-Centre Cologne, Germany (https://mpgc.mpipz.mpg.de/home/). rRNA depletion was conducted with 1 μg total RNA using Ribo-Zero rRNA Removal Kit Bacteria (Illumina), followed by library preparation with NEBNext Ultra Directional RNA Library Prep Kit for Illumina (New England Biolabs). Library preparation included 11 cycles of PCR amplification. Quality and quantity were assessed at all steps via capillary electrophoresis (TapeStation, Agilent Technologies) and fluorometry (Qubit, Thermo Fisher Scientific). Sequencing was performed on HiSeq 3000 (Illumina) with 1× 150 bp single reads. Libraries were re-sequenced until a sufficient number of reads were obtained. Sequencing files can be downloaded from EBI ArrayExpress under accession number E-MTAB-11043.

### Cappable-sequencing

Total RNA was isolated from *M. xanthus* cells developed under submerged conditions as described. Library preparation and sequencing was performed at Vertis Biotechnologie AG, Freising, Germany (https://www.vertis-biotech.com/home) as described in (62). Briefly, 5’ triphosphorylated RNA was capped with 3’-desthiobiotin-TEG-guanosine 5’ triphosphate (DTBGTP) (New England Biolabs) using the vaccinia capping enzyme (VCE) (New England Biolabs). Then biotinylated RNA molecules were captured using streptavidin beads and eluted with a biotin-containing buffer. RNA samples were poly(A)-tailed using poly(A) polymerase. Then the 5’-PPP or CAP structures were converted to 5’-P using CAP-Clip Acid Pyrophosphatase (Cellscript). Afterwards, an RNA adapter was ligated to the newly formed 5’-monophosphate structures. First-strand cDNA synthesis was performed using an oligo(dT)-adapter primer and M-MLV reverse transcriptase. The resulting cDNAs were PCR-amplified using a proof-reading enzyme. The libraries were amplified in 15 cycles of PCR. The generated cDNA libraries were sequenced on an Illumina NextSeq 500 system using 75 bp read length. Sequencing files can be downloaded at EBI ArrayExpress under accession number E-MTAB-11042.

### Organism

The genome and annotation of *Myxococcus xanthus* DK 1622 (NC_008095.1, downloaded 28.01.2019) were used for all analyses.

### RNA-seq analysis

All sequencing runs of one sample were concatenated using “cat” (GNU coreutils 8.30). As reverse transcription is part of the sequencing protocol, this was compensated for by “reverse_complement” of the FASTX-Toolkit 0.0.14 (http://hannonlab.cshl.edu/fastx_toolkit). The differential gene expression analysis was done using the RNA-seq pipeline Curare 0.2.1. This software will be described in details in a separate manuscript. Briefly, the reads were aligned using Bowtie2 2.4.2 in ‘very-sensitive’ mode and with ‘--mm’ option (75). Except for the WT_t24_2 sample, all samples had mapping rates higher than 90% (Table S5). The resulting SAM/BAM files were processed with Samtools 1.12 (76). The subsequent assignment of mapped reads to genome features was done using the featureCounts (77) of the subread 2.0.1 package (78). featureCounts was run with “-s 1” settings assigning reads strand specific to the ‘gene’ features. For every sample, more than 93% of all reads could be assigned to a ‘gene’ feature (Table S6). Finally, the differential gene expression was analyzed with DESeq2 1.30.1 (79). The Curare version of this analysis can be downloaded at Zenodo (DOI: 10.5281/zenodo.5541852). The count table and mapping results can be downloaded from EBI ArrayExpress under accession number E-MTAB-11043.

### Cappable-seq analysis

The TSS pipeline in (62) was used for TSS detection with modifications. This modified pipeline will be described in detail in a separate manuscript. Briefly, the raw Cappable-seq reads were mapped with Bowtie2 2.4.1 using ‘--all’, ‘--mm’, and ‘--very-sensitive’ settings (75). As in the RNAseq analysis, all samples except WT_t24_2 had a mapping rate of >90% (Table S7). A custom script was used to filter all non-best mappings of each read (two equal good mappings will be counted as half a read/mapping each). Created SAM and BAM files were processed using Samtools 1.12 (76) and Pysam 0.16 (https://pysam.readthedocs.io/). Only the first base of each mapping was used for building’ alignments per base’ scores (Rns) and every following step. The following formula, altered from (62), was used to normalize these scores: RRS = (Rns/Rt) * 1,000,000 (RRS: relative read score, Rt: total number of reads mapped). As in (62), an RRS of 1.5 was used as the lower threshold. The first mapped nucleotide from the sequencing reads identifies the orientation and position of the first nucleotide of the primary transcript. TSSs within three nucleotides were clustered into one TSS. In case of flanking clusters or TSSs within a distance of three or less nucleotides, these were merged into one large cluster. The TSS with the highest RRS in a cluster was defined as the major TSS and used in these analyses. The complete pipeline can be downloaded at Zenodo (DOI: 10.5281/zenodo.5541852). The mapping and TSS results can be downloaded from EBI ArrayExpress under accession number E-MTAB-11042.

### RT-qPCR

1 μg of total RNA isolated as described above was used to synthesize cDNA with the High Capacity cDNA Reverse Transcription Kit (Applied Biosystems) according to the manufacturer’s protocol. cDNA templates were diluted 10-fold, 2 μl of a diluted sample was used as a template for RT-qPCR reaction, which contained 1× SYBR Green PCR Master Mix (Applied Biosystems), 2.5 μM of each primer and H_2_O to a final volume of 25 μl. A 7500 Real time PCR detection system (Applied Biosystems) was used for RT-qPCR measurements using standard conditions. Experiments were done in two biological replicates, each in two technical replicates. Relative gene expression levels were calculated using the comparative Ct method.

### Operon mapping

RNA preparation was done as described. Primers used are listed in Table S4, and used on genomic DNA, RNA without addition of reverse transcriptase, and cDNA.

### Immuno-blot analysis

Immunoblots were carried out as described (74). Rabbit polyclonal α-DmxB (1:1000 dilution) (40), α-GFP (Roche, 1:2000 dilution), α-mCherry (Biovision, 1:2000 dilution), α-protein C (1:2000 dilution) (80) and α-PilC (1:5000 dilution) (81) antibodies were used together with horseradish-conjugated goat anti-rabbit immunoglobulin G (Sigma-Aldrich) or anti-mouse sheep IgG antibody (GE Healthcare) as secondary antibody. Blots were developed using Luminata crescendo Western HRP Substrate (Millipore) and visualized using a LAS-4000 luminescent image analyzer (Fujifilm). To quantify immuno-blots, signal intensities of the relevant protein bands were quantified using Fiji (82) and normalized relative to the PilC loading control from the same blot. All immuno-blots were performed in three independent biological replicates.

### Protein purification

To purify His_6_-MrpC, *E. coli* Rosseta 2 (DE3)/pLysS strain (Novagen) was transformed with pPH158 (33). The culture was grown in 1L LB with addition of chloramphenicol and kanamycin at 37ºC to an optical density at 600 nm of 0.5-0.7. Protein expression was induced by addition of isopropylthio-β-galactoside (IPTG) to a final concentration of 0.5 mM for 3 h at 37ºC. Cells were harvested by centrifugation at 5,000*g* for 10 min at 4ºC and resuspended in lysis buffer (50 mM NaH_2_PO_4_, 300 mM NaCl, 5 mM MgCl_2,_ 10 mM imidazole, 5% glycerol, pH 8.0 and Complete Protease Inhibitor Cocktail Tablet (Roche)). Cells were disrupted using a French press and harvested at 48,000*g* for 40 min at 4ºC. The cleared supernatant was filter with 0.45 μm sterile filter (Millipore Merck, Schwalbach) and applied to column with 2 ml of Ni^2+^-NTA-agarose (GE Healthcare) equilibrated with wash buffer (50 mM NaH_2_PO_4_, 300 mM NaCl, 5 mM MgCl_2_, 50 mM imidazole, 5% glycerol, pH 8.0). Protein was eluted with elution buffer (50 mM NaH_2_PO_4_, 300 mM NaCl, 5 mM MgCl_2_, 100-500 mM imidazole, 5% glycerol, pH 8.0). Fractions containing purified His_6_-MrpC were combined and loaded onto a HiLoad 16/600 Superdex 200 pg (GE Healthcare) size exclusion chromatography column equilibrated with lysis buffer without imidazole. Fractions containing His_6_-tagged MrpC were frozen in liquid nitrogen and stored at −80°C.

### Electrophoretic mobility shift assay (EMSA)

Hex-labelled probes were generated using the primer pairs listed in Table S4 and plasmids containing the WT or mutant promoters as templates. Assays were performed as described (83). Briefly, purified His_6_-MrpC was mixed at the indicated concentrations with 6 nM (*dmxB* fragments) or 10 nM (*pmxA* fragments) of HEX-labeled DNA fragment in reaction buffer (10 mM Tris pH 8.0, 50 mM KCl, 1 mM DTT, 10 μg ml^-1^ BSA, 10% glycerol, 0.5 μg herring sperm DNA (Thermo Fisher Scientific)) in a total volume of 10 μl, and incubated for 15 min at 20°C. Reaction samples were separated on a 5% polyacrylamide gel in 0.5× TBE (45 mM Tris, 45 mM Borate, 1 mM EDTA) for 1.5 h. Gels were imaged using a Typhoon Phosphoimager (GE Healthcare).

### c-di-GMP and cGAMP quantification

To quantify the c-di-GMP and cGAMP levels, cells were grown in CTT or developed under submerged conditions as described. Cells were harvested at 2,500*g* for 20 min at 4°C, lysed in extraction buffer (HPLC grade acetonitrile/methanol/water (2/2/1, v/v/v)), and supernatants evaporated to dryness in a vacuum centrifuge. Pellets were dissolved in HPLC grade water and analyzed by LC-MS/MS. c-di-GMP and cGAMP quantification was performed at the Research Service Centre Metabolomics at the Hannover Medical School, Germany. Experiments were done in three biological replicates. Protein concentrations were determined in parallel using a Pierce^®^Microplate BCA Protein Assay Kit (Thermo Scientific).

### Bioinformatics

Heatmaps were created using R package pheatmap (https://cran.r-project.org/web/packages/pheatmap/index.html). Protein domains were identified using Pfam v33.1 (pfam.xfam.org) (84); signal peptides were predicted with SignalP 5.0 (www.cbs.dtu.dk/services/SignalP-5.0) (85).

