## Supplementary Information for "The CRP-like transcriptional regulator MrpC curbs c-di-GMP and 3’, 3’ cGAMP nucleotide levels during development in *Myxococcus xanthus*"

This file contains:

- Supplementary figures 1-7
- Supplementary Materials and Methods.
- Supplementary Tables 4-7

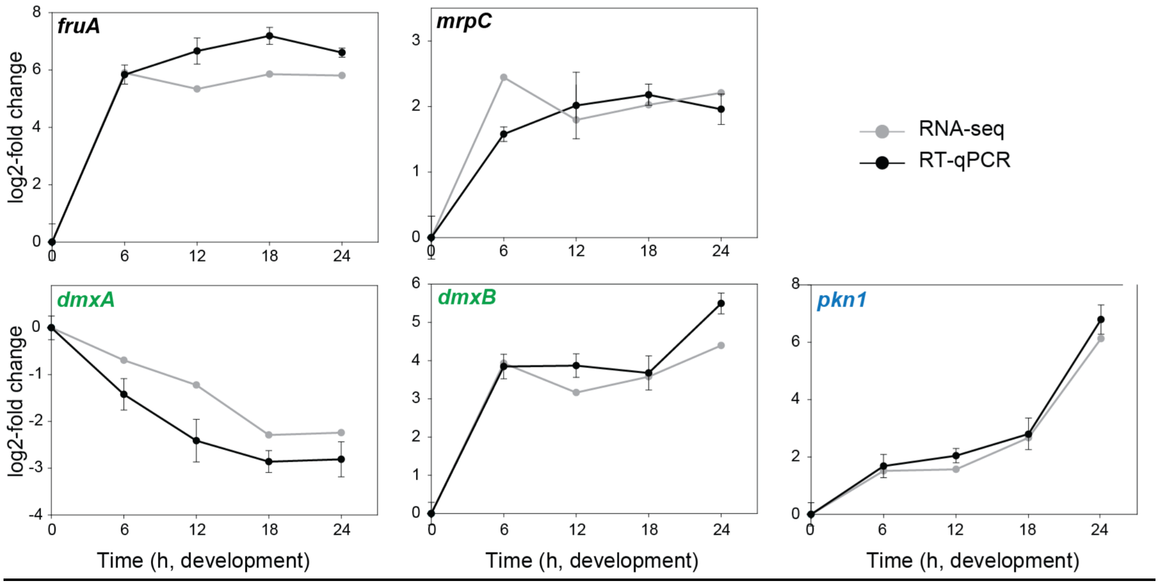


Supplementary Figure 1. Benchmarking of RNA-seq expression data using RT-qPCR
Total RNA was isolated from WT cells developed in MC7 submerged cultures at the indicated time points. Both datasets are shown as log2-fold change at 6, 12, 18 or 24 h of development compared to 0 h. RT-qPCR data are shown as mean±SD from two biological replicates, each with two technical replicates, relative to 0 h. The same RNA samples were used in RNA-seq and RT-qPCR analyses.

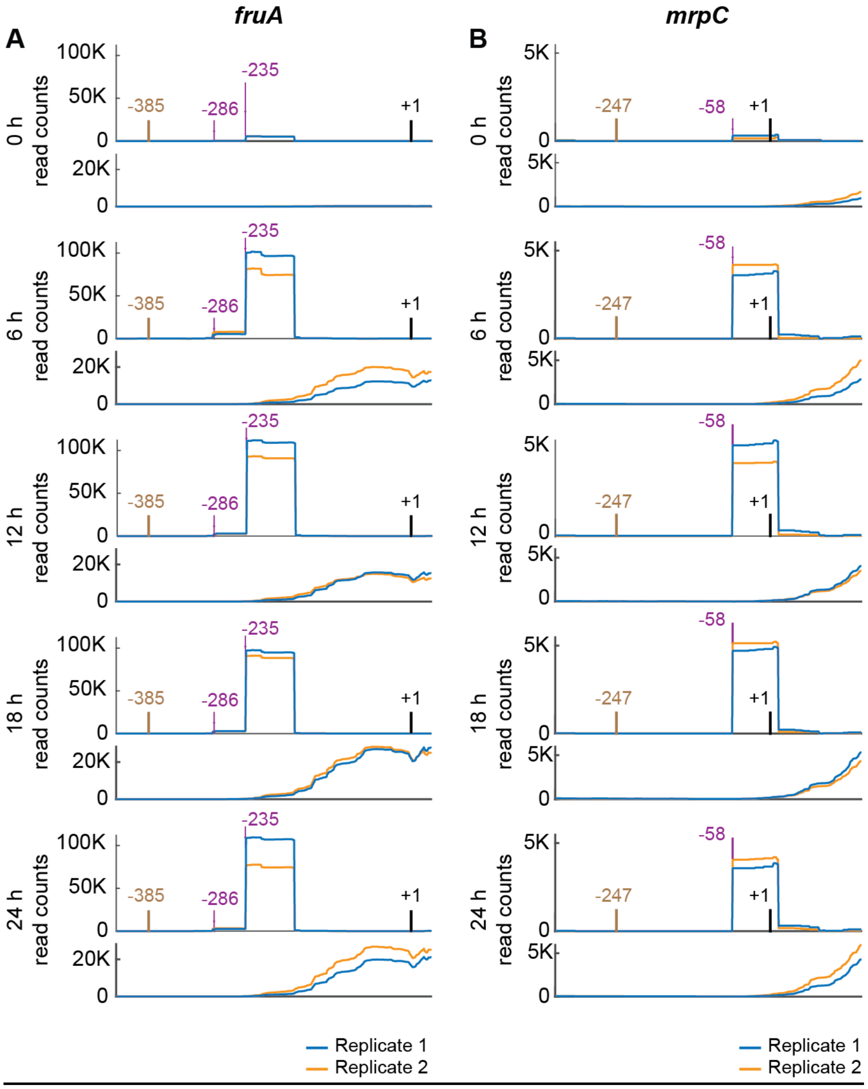


Supplementary Figure 2. Benchmarking of transcriptional start site mapping using Cappable-seq

A, B. Analysis of *fruA* and *mrpC* promoter regions with Cappable-seq and RNA-seq. RNA-seq (lower panels) and Cappable-seq (upper panels) data are visualized at different time points. For each time point, mapped read counts for both biological replicates are shown in blue and orange. The data from RNA-seq and Cappable-seq were obtained from different samples. +1 indicates the TSC. TSSs are indicated in purple relative to the TSC. The center of the MrpC ChIP-seq peak is in brown. Only the highest scoring TSSs are included. For all potential TSSs, see Table S3A.

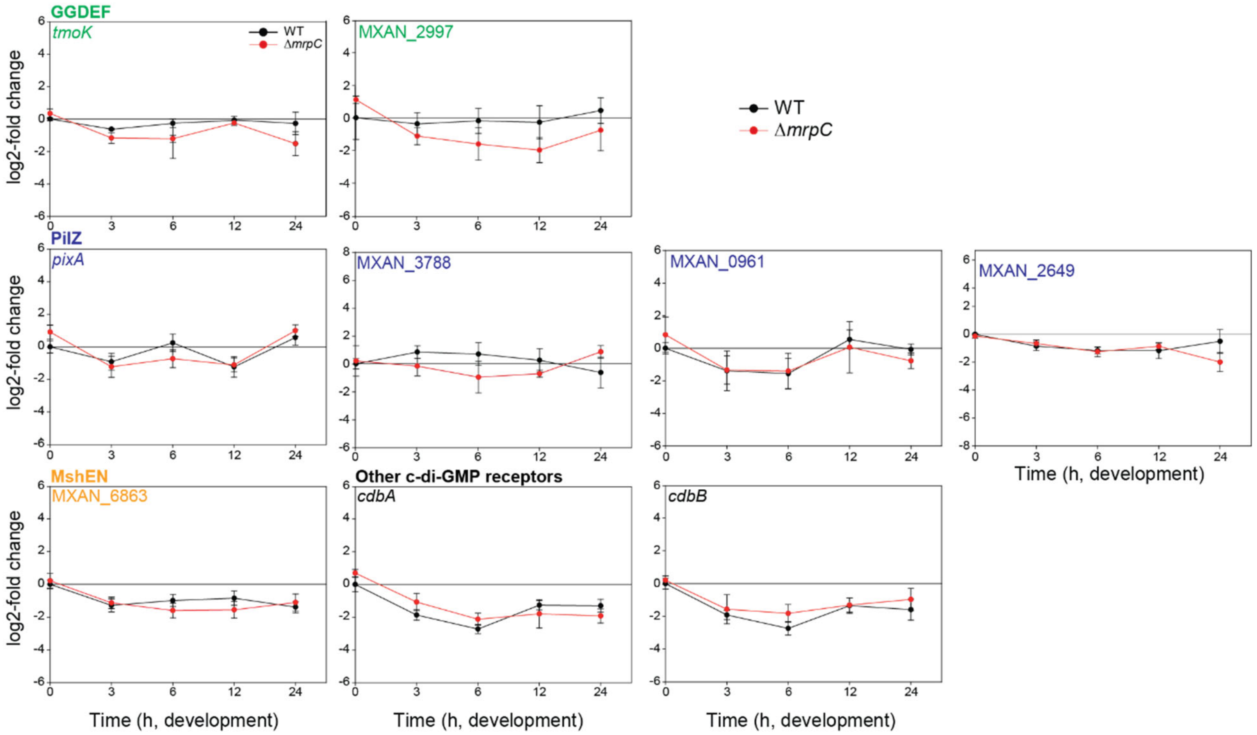


Supplementary Figure 3. Expression of genes encoding "c-di-GMP associated proteins" potentially regulated by MrpC

Total RNA was isolated from cells developed in MC7 submerged cultures at the indicated time points of WT (black) and the Δ*mrpC* mutant (red). Transcript levels are shown as mean±SD from two biological replicates, each with two technical replicate, relative to WT at 0 h.

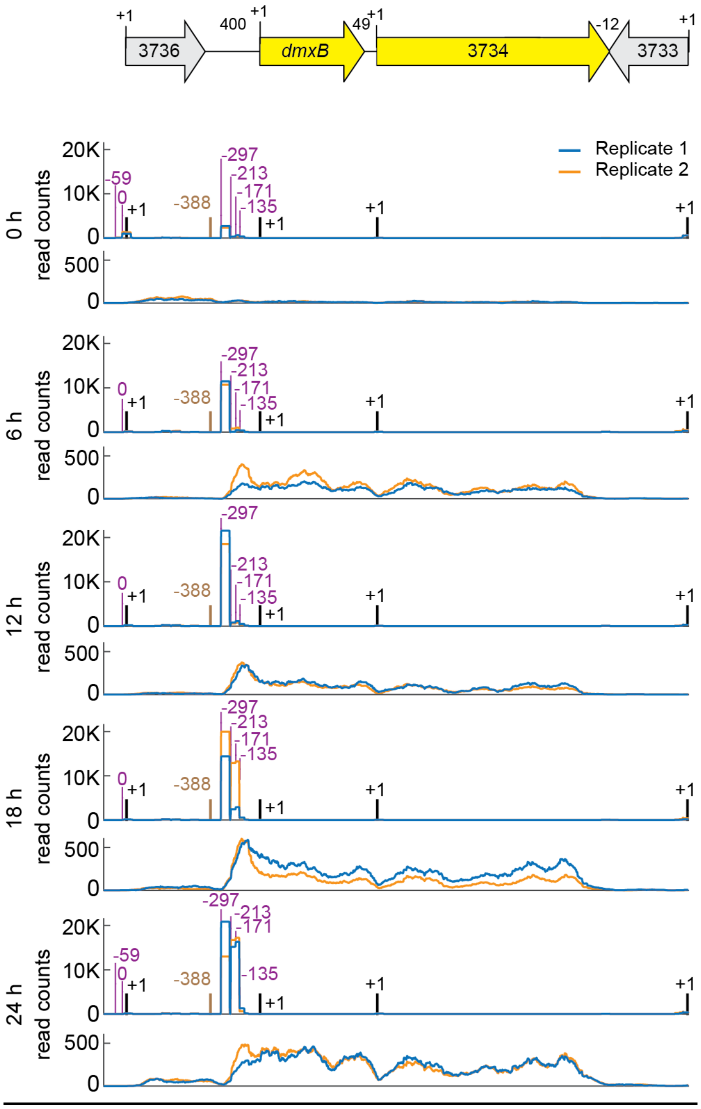


Supplementary Figure 4. Mapping of the *dmxB* locus with RNA-seq and Cappable-seq.
Upper part, *dmxB* locus with numbers above indicating distance between start and stop codons. Lower part, visualization of RNA-seq (lower panels) and Cappable-seq (upper panels) data at different time points. For each time point, mapped read counts for both biological replicates are shown in blue and orange. The data from RNA-seq and Cappable-seq were obtained from different samples. +1 indicates TSC of gene as shown in upper part. Potential TSSs are indicated in purple relative to the relevant TSC. The center of the MrpC ChIP-seq peak is in brown.

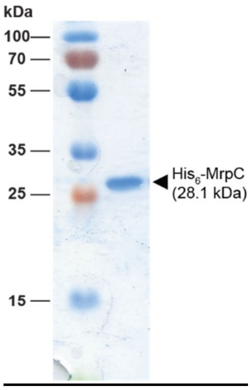


Supplementary Figure 5. Purification of His$_6$-MrpC.
Purified His$_6$-MrpC protein was separated by SDS-PAGE and stained with Coomassie blue.

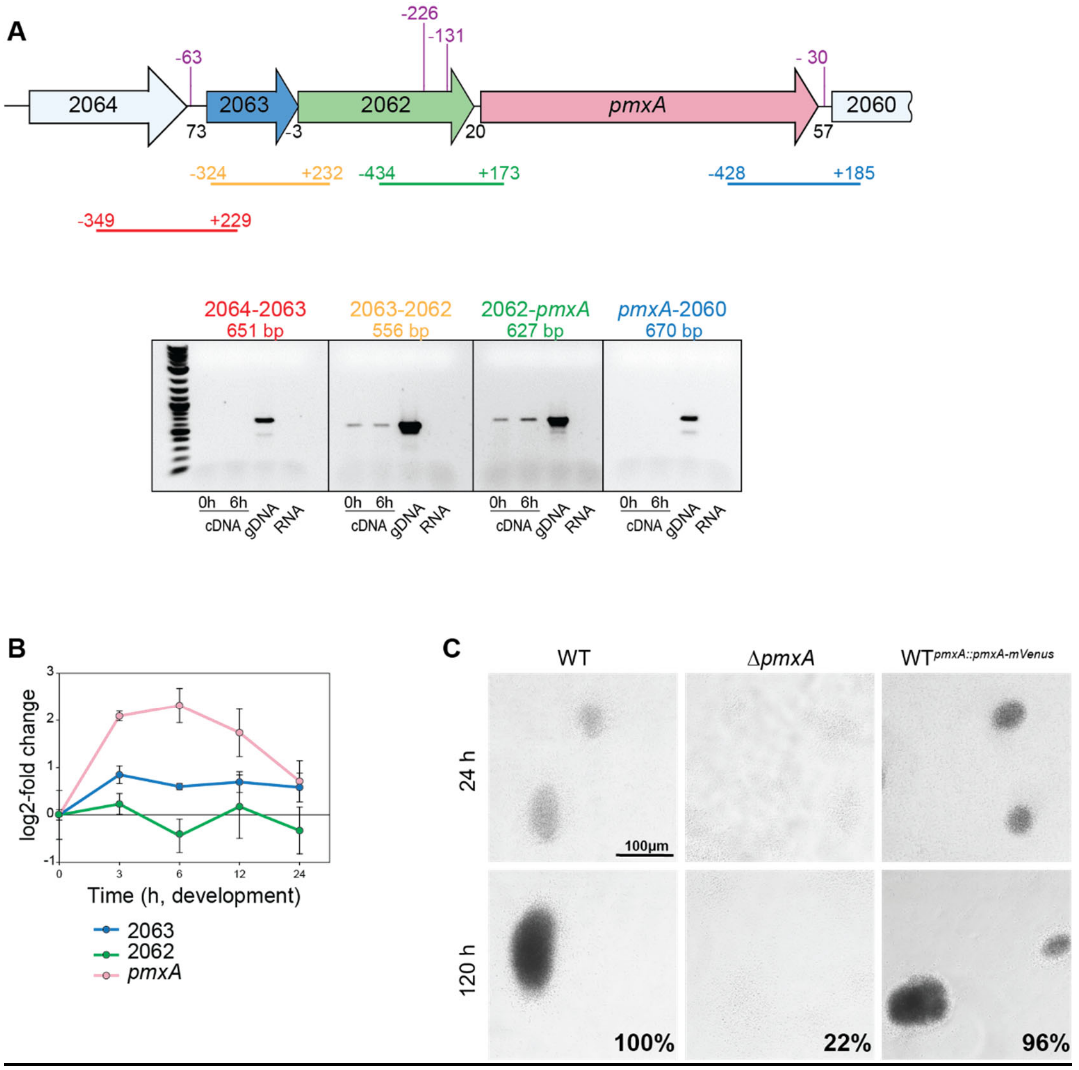


Supplementary Figure 6. Operon mapping of the *pmxA* locus

A. Schematic of *pmxA* locus and operon mapping. Upper panel, *pmxA* locus in which the direction of transcription is indicated by the arrows. Numbers below indicate distance between start and stop codons of flanking genes; numbers above in purple indicate potential TSSs as mapped by Cappable-seq relatively to the nearest TSC. Coloured bars below indicate the fragments amplified in the RT-qPCR analysis; coordinates indicate the 5'- and 3'-ends of the amplified fragment relative to the relevant start and stop codon. Lower panel, agarose gel with results from operon mapping. The fragments indicated in red, orange, green and blue were amplified from cDNA prepared from total RNA of vegetative (0 h) and developing (6 h) cells, from genomic DNA (gDNA) or from total RNA without addition of reverse transcriptase (RNA). Primer pairs used: SK318 and SK319 (fragment 2060-*pmxA*), SK320 and SK321 (fragment *pmxA*-2062), SK322 and SK323 (fragment 2062-2063) and SK324 and SK325 (fragment 2063-2064).

B. Comparison of *pmxA*, MXAN_2063 and MXAN_2062 expression during development. Total RNA was extracted at the indicated points from WT cells developed under MC7 submerged conditions. Transcripts levels are shown as log2-fold changes relative to 0 h as mean±SD from two biological replicates, each in two technical replicate.

C. Developmental assays for WT, ∆*pmxA* mutant and WT producing PmxA-mVenus from the native site. Development was performed under MC7 submerged conditions and cells imaged at 24 and 120 h. Numbers indicate heat- and sonication resistant spores at 120 h as percentage of WT (100%). Scale bar, 100μm.

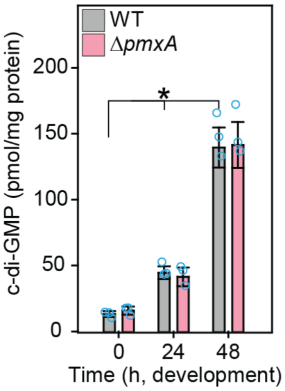


Supplementary Figure 7. c-di-GMP accumulation in WT and Δ*pmxA* mutant.

Cells were harvested at the indicated time points of development, and nucleotide levels and protein concentrations determined. Levels are shown as mean±SD calculated from three biological replicates. Individual data points are in light blue. *, *P*-value <0.05 in Student's t-test.

**Supplementary Materials and Methods.**

**Plasmid construction**

Plasmid **pSK29** was used to integrated *pmxA*-mVenus into native site in genome. The upstream and downstream regions of the *pmxA* were amplified using primer pairs SK44-SK-083 and SK084-SK47. Obtained fragments were fused during second round of PCR amplification. The product was cloned into KpnI/XbaI sites of pBJ114.

Plasmid **pSK65** was used to integrate mCherry copy into WT. To amplify mCherry primers mCherry_F_EcoRI - mCherry _R_XbaI were used. The product was cloned into EcoRI/XbaI sites of pSWU30.

Plasmid **pSK81** was used to create mCherry reporter for *pmxA* promoter. To amplify *pmxA* promoter fused to mCherry SK278 - SK277 and SK276 - mCherry_R_XbaI primers were used. Obtained fragments were fused during second round of PCR amplification using SK278 - mCherry_R_XbaI primers.The product was cloned into EcoRI/XbaI sites of pSWU30.

Plasmid **pSK103** was used to create mCherry reporter for *pmxA* promoter with mutated BS1.To introduce mutations in BS1 of *pmxA* promoter, SK278 - SK301 and SK300 - mCherry_R_XbaI primers were used on **pSK81** as a template. Obtained fragments were fused during second round of PCR amplification using SK278 - mCherry_R_XbaI primers.The product was cloned into EcoRI/XbaI sites of pSWU30.

Plasmid **pSK105** was used to create mCherry reporter for *pmxA* promoter with mutated BS2.To introduce mutations in BS2 of *pmxA* promoter, SK278 - SK309 and SK308 - mCherry_R_XbaI primers were used on **pSK81** as a template. Obtained fragments were fused during second round of PCR amplification using SK278 - mCherry_R_XbaI primers.The product was cloned into EcoRI/XbaI sites of pSWU30.

Plasmid **pSK114** was used to create mCherry reporter for *pmxA* promoter with mutated BS3.To introduce mutations in BS3 of *pmxA* promoter, SK278 - SK346 and SK345 - mCherry_R_XbaI primers were used on **pSK81** as a template. Obtained fragments were fused during second round of PCR amplification using SK278 - mCherry_R_XbaI primers.The product was cloned into EcoRI/XbaI sites of pSWU30.

Plasmid **pSK111** was used to create mCherry reporter for *pmxA* promoter with mutated BS1 and BS2. To introduce mutations in BS1 of *pmxA* promoter, SK278 - SK301 and SK300 - mCherry_R_XbaI primers were used on **pSK105** as a template. Obtained fragments were fused during second round of PCR amplification using SK278 - mCherry_R_XbaI primers.The product was cloned into EcoRI/XbaI sites of pSWU30.

Plasmids **pSK101** was used to create mCherry reporter for *dmxB* promoter. To amplify *dmxB* promoter fused to mCherry SK279-SK280 and SK281 - mCherry_R_XbaI primers were used. Obtained fragments were fused during second round of PCR amplification using SK279 - mCherry_R_XbaI primers.The product was cloned into EcoRI/XbaI sites of pSWU30.

Plasmids **pSK109** was used to create mCherry reporter for *dmxB* promoter with mutated BS4. To introduce mutations in BS4 of *dmxB* promoter, SK279 - SK317 and SK316 - mCherry_R_XbaI primers were used on **pSK101** as a template. Obtained fragments were fused during second round of PCR amplification using SK279 - mCherry_R_XbaI primers.The product was cloned into EcoRI/XbaI sites of pSWU30.

Plasmids **pSK112** was used to create mCherry reporter for *dmxB* promoter with mutated BS3. To introduce mutations in BS3 of *dmxB* promoter, SK279 - SK305 and SK304 - mCherry_R_XbaI primers were used on **pSK101** as a template. Obtained fragments were fused during second round of PCR amplification using SK279 - mCherry_R_XbaI primers.The product was cloned into EcoRI/XbaI sites of pSWU30.

Plasmids **pSK115** was used to create mCherry reporter for *dmxB* promoter with mutated BS2. To introduce mutations in BS2 of *dmxB* promoter, SK279 - SK345 and SK344 - mCherry_R_XbaI primers were used on **pSK101** as a template. Obtained fragments were fused during second round of PCR amplification using SK279 - mCherry_R_XbaI primers.The product was cloned into EcoRI/XbaI sites of pSWU30.

Plasmids **pSK121** was used to create mCherry reporter for *dmxB* promoter with mutated BS1. To introduce mutations in BS2 of *dmxB* promoter, SK279 - SK356 and SK355 - mCherry_R_XbaI primers were used on **pSK101** as a template. Obtained fragments were fused during second round of PCR amplification using SK279 - mCherry_R_XbaI primers.The product was cloned into EcoRI/XbaI sites of pSWU30.

**Supplementary Tables.**

**Table S1.** RNA-seq analysis of *M. xanthus* genes encoding c-di-GMP associated proteins in WT (Excel file).

**Table S2.** Transcriptional Start Sites (TSSs) with RRS 1.5 cutoff in *M. xanthus* genome (Excel file).

**Table S3.** Combined analysis of Transcriptional Start Sites (TSSs) and MrpC ChIPseq data for *fruA* and *mrpC* (A), genes encoding c-di-GMP associated proteins (B), and their operons (C) (Excel file).

**Table S4.** Primers used in this study

| Name | Sequence (5'→3') | Description |
|---|---|---|
| SK44 | GCGCGGTACCAGATGAAGCTCACCGAGC | Endogenous substitution of *pmxA* to *pmxA*-mVenus |
| SK47 | GCGCTCTAGAGGCCGCTCGAATCCCTCCG | |
| SK83 | CTTCGGCCGTTACTTGTACAGCTCGTC | |
| SK84 | TACAAGTAACGGCCGAAGGATGGTAGG | |
| mCherry_R XbaI | ATCGTCTAGATTACTTGTACAGCTCGTCCATGCC | mCherry |
| SK279 | CGGCGAATTCTCCGCCGCCGGTCATGCC | Cloning of P*dmxB* mCherry |
| SK280 | AATGAAGCGGAGTGGTGCATGGTGAGCAAGGGCGAG | |
| SK281 | CTCGCCCTTGCTCACCATGCACCACTCCGCTTCATT | |
| SK355 | GTTGTGGGCGTTGGG**TAAG**GCTTGCAGGACCGGG | P*dmxB* BS1 site directed mutagenesis |
| SK356 | CCCGGTCCTGCAAGC**CTT**ACCCAACGCCCACAAC | |
| SK343 | AGGGGCGTTGGGAGC**CTTA**GGAAGGGCGTGGCAC | P*dmxB* BS2 site directed mutagenesis |
| SK344 | GTGCCACGCCCTTCC**TAAG**GCTCCCAACGCCCCT | |
| SK304 | CGCGCGGCCACCGAG**GAAT**CGCCCGCGCACGGGAATATCTAC | P*dmxB* BS3 site directed mutagenesis |
| SK305 | GTAGATATTCCCGTGCGCGGGCG**ATTC**CTCGGTGGCCGCGCG | |
| SK316 | TGCAGTTCAGCACCC**ATTC**CTAGCGACCCGTCGC | P*dmxB* BS4 site directed mutagenesis |
| SK317 | GCGACGGGTCGCTAG**GAAT**GGGTGCTGAACTGCA | |
| SK276 | CCCTGGAGGAGCCACGCCATGGTGAGCAAGGGCGAG | Cloning of P*pmxA* mCherry |
| SK277 | CTCGCCCTTGCTCACCATGGCGTGGCTCCTCCAGGG | |
| SK278 | CGGCGAATTCTGAAGCGACGCCCCGACCGG | |
| SK300 | CAGGAGCAACGCGC**GAAT**TTCACGCAGACAGTC | P*pmxA* BS1 site directed mutagenesis |
| SK301 | GACTGTCTGCGTGAA**ATTC**GCGCGTTGCTCCTG | |
| SK308 | TACGGGGCGCGGAA**GAAT**ATGAAGTGGAAGTGT | P*pmxA* BS2 site directed mutagenesis |
| SK309 | ACACTTCCACTTCAT**ATTC**TTCCGCGCCCCGTA | |
| SK345 | CGACGCCCCGACCGG**ATTC**GGCACCTGGGCCCGG | P*pmxA* BS3 site directed mutagenesis |
| SK346 | CCGGGCCCAGGTGCC**GAAT**CCGGTCGGGGCGTCG | |
| 4361_qPCR_forw | CACCGACAAGCGCAAGCAG | RT-qPCR, *cdbA* |
| 4361_qPCR_rev | GCACGACCCAGGAAAGGGA | |
| 4362_qPCR_forw | CCGAGGACATGCTGGAGGAG | RT-qPCR, *cdbB* |
| 4362_qPCR_rev | CGTTGACCGACGGCATCTTC | |
| qdmxAf | GTTTCGAGGACAACAACGGG | RT-qPCR, *dmxA* |
| qdmxAr | AAGATCTTCAGCGCACCCAG | |
| q3735f | GGTCCCTTCTGCTCATCATC | RT-qPCR, *dmxB* |
| q3735r | AGGAACCTGTCCAGGAGGA | |
| fruA for | ATCATCTCGCAGTGCTTCGA | RT-qPCR, *fruA* |
| fruA rev | CCTCGGACCAGGGAGTTGA | |

| | | |
|---|---|---|
| qmrpcCf | CCGCACAACTCGACCATCTA | RT-qPCR, *mrpC* |
| qmrpcCr | GGTGCTGTTCTTGCCGATG | |
| 7440 qPCR forw | GTGGACTTCCTCTGCGAATC | RT-qPCR, *nla24* |
| 7440 qPCR rev | GATGACCAGGTCGAAGGACT | |
| q1087f | AGCTGCACGGAAAGGTCCCC | RT-qPCR, *pixA* |
| q1087r | AGGGCGAAGCCGTCCATCAC | |
| q1467f | CCCTGTCGGTCATGTGTGAA | RT-qPCR, *pkn1* |
| q1467r | ACGAACTGGACGCCGAAG | |
| pmxA qPCR fw | GGTGGACATCAAGCAGAAGA | RT-qPCR, *pmxA* |
| pmxA qPCR rev | CCTTCTCCAGGCTCTCGTAG | |
| q4445f | ACCGCATCATCCCGCTTTAT | RT-qPCR, *tmoK* |
| q4445r | ACACGCTCATGATGGGGAA | |
| q0961f | TGGTGATGTGGGCTGCTGGG | RT-qPCR, MXAN_0961 |
| q0961r | CCAAGGCGGAGCGAGATGCT | |
| q1525f | CACCAACGGTACCTTCCTCAA | RT-qPCR, MXAN_1525 |
| q1525r | GATGGTCTCGTGGTACTGGG | |
| q2649f | TCACGCCGTTGATTGAGATGA | RT-qPCR, MXAN_2649 |
| q2649r | GCAAGTAGAAGGTGCTCTCCA | |
| q2902f | TGGACGAAATCGAGAACACCA | RT-qPCR, MXAN_2902 |
| q2902r | TGAGCCGGTAGAAGAGGTCC | |
| q2997f | TCAAGGACGAGTTGGAGGAC | RT-qPCR, MXAN_2997 |
| q2997r | ATGGAGAACTCGGGGATGC | |
| q3788f | GGCTTGTCGGTGTTGATGTA | RT-qPCR, MXAN_3788 |
| q3788r | AACACCTCGACCCCTCATCC | |
| q4232f | TGCACGACATCGGGAAGATT | RT-qPCR, MXAN_4232 |
| q4232r | CGGAATGGCCTGGATCATCT | |
| q6863f | AGAGCAAGCAGAAGCGCGGA | RT-qPCR, MXAN_6863 |
| q6863r | CCAGCTCCTCGACGCGATCC | |
| q6957f | GTGTGGTCATCGCATCCTCT | RT-qPCR, MXAN_6957 |
| q6957r | CGCCTTCCCGAGACAACA | |
| q7024f | TCTTCACGGTGGTGGAGTTC | RT-qPCR, MXAN_7024 |
| q7024r | ACACCTTCAGCGTCAGCC | |
| q7500f | CAGCTCTACCGGGGTGAAAC | RT-qPCR, MXAN_7500 |
| q7500r | GGAGCCTCCATGTTCGTCAG | |
| SK318 | CGCTTGTTCGTCCATTCGTC | *pmxA* operon mapping |
| SK319 | GGGAAGATTGGCATCGTGGA | |
| SK320 | CCGAGGAAGATTTCGGCCTT | |
| SK342 | GTCCGCGTGGAGGCCGAG | |

| | | |
|---|---|---|
| SK322 | CCGTCTCCAGGGCCTTCTGC | |
| SK323 | GGGCGGCGTCCAAATCAAGC | |
| SK324 | TGTCATTCTCGTAGAGCGGC | |
| SK325 | TCTGCTACGACCTGCGATTC | |
| ePmxA-HEX_F | TGAAGCGACGCCCCGACC | Amplification of Hex-labeled P*pmxA* EMSA probe |
| SK351_ R | GGCGTGGCTCCTCCAGGGGT | |
| eDmxB-HEX_F | TCCGCCGCCGGTCATGCC | Amplification of Hex-labeled P*dmxB* EMSA probe |
| SK352_R | GCACCACTCCGCTTCATTCG | |

* Restriction sites are underlined and mutations introduced by site-directed mutagenesis are in bold.

**Table S5.** Mapping rates for RNA-seg[1]

| Sample | #Reads | Aligned 0 times | Aligned 1 times | Aligned >1 times |
|---|---|---|---|---|
| wt_t0_1 | 16,338,933 | 734,180 | 15,070,943 | 533,810 |
| wt_t0_2 | 20,997,454 | 246,610 | 20,146,141 | 604,703 |
| wt_t6_1 | 14,199,176 | 794,742 | 12,924,338 | 480,096 |
| wt_t6_2 | 21,155,374 | 1,038,091 | 19,318,904 | 798,379 |
| wt_t12_1 | 17,345,707 | 1,000,489 | 16,202,341 | 142,877 |
| wt_t12_2 | 20,957,831 | 586,779 | 19,974,379 | 396,673 |
| wt_t18_1 | 19,298,934 | 1,008,536 | 18,075,039 | 215,359 |
| wt_t18_2 | 20,855,167 | 662,083 | 19,469,907 | 723,177 |
| wt_t24_1 | 17,719,468 | 1,166,915 | 16,373,886 | 178,667 |
| wt_t24_2 | 21,138,562 | 2,402,535 | 18,191,387 | 544,640 |

[1] Bowtie2 (Single End, --very-sensitive, --mm, version 2.4.2) mapping statistics of the *Myxococcus xanthus* DK 1622 differential gene expression experiment. NC_008095.1 (downloaded 28.01.2019) was used as the reference genome.

**Table S6.** Feature assignment of RNA-seg data[1]

| Sample | #Reads | Assigned | Unassigned (No Feature) | Unassigned (Ambiguity) |
|---|---|---|---|---|
| wt_t0_1 | 15,604,753 | 14,572,563 | 144,854 | 887,336 |
| wt_t0_2 | 20,750,844 | 19,356,757 | 231,753 | 1,162,334 |
| wt_t6_1 | 13,404,434 | 12,633,493 | 112,716 | 658,225 |
| wt_t6_2 | 20,117,283 | 19,037,858 | 188,655 | 890,770 |
| wt_t12_1 | 16,345,218 | 15,358,948 | 149,592 | 836,678 |
| wt_t12_2 | 20,371,052 | 19,346,107 | 174,754 | 850,191 |
| wt_t18_1 | 18,290,398 | 17,085,203 | 161,137 | 1,044,058 |
| wt_t18_2 | 20,193,084 | 19,140,220 | 164,206 | 888,658 |
| wt_t24_1 | 16,552,553 | 15,460,279 | 147,983 | 944,291 |
| wt_t24_2 | 18,736,027 | 17,505,669 | 182,409 | 1,047,949 |

[1] FeatureCounts (Assigned to 'gene' features, Subread version 2.0.1) statistics of the *Myxococcus xanthus* DK 1622 differential gene expression experiment. The NC_008095.1 annotation was used. #Reads: Total number of reads used in FeatureCounts; Assigned: Number of reads assigned to a 'gene' feature. Unassigned (No Feature): Reads, which were aligned to areas without any features; Unassigned (Ambiguity): Reads, which could be assigned to at least two features.

**Table S7.** Mapping rates for Cappable-seq samples [1]

| Sample | #Reads | Aligned 0 times | Aligned 1 times | Aligned >1 times |
|---|---|---|---|---|
| wt_t0_1 | 12,257,955 | 501,139 | 10,712,553 | 1,044,263 |
| wt_t0_2 | 11,807,934 | 437,614 | 10,561,172 | 809,148 |
| wt_t6_1 | 12,852,056 | 693,624 | 10,701,275 | 1,457,157 |
| wt_t6_2 | 13,170,210 | 560,070 | 11,710,213 | 899,927 |
| wt_t12_1 | 13,315,318 | 836,975 | 9,680,933 | 2,797,410 |
| wt_t12_2 | 11,899,311 | 756,439 | 8,652,199 | 2,490,673 |
| wt_t18_1 | 11,971,162 | 763,512 | 8,285,279 | 2,922,371 |
| wt_t18_2 | 12,536,870 | 1,239,561 | 9,200,524 | 2,096,785 |
| wt_t24_1 | 12,388,364 | 841,226 | 9,388,384 | 2,158,754 |
| wt_t24_2 | 12,775,299 | 1,981,621 | 9,133,926 | 1,659,752 |

[1] Bowtie2 (Single End, --very-sensitive, --mm, --all, version 2.4.1) mapping statistics of the *Myxococcus xanthus* DK 1622 differential gene expression experiment. NC_008095.1 (downloaded 28.01.2019) was used as the reference genome.
